# Small molecule stabilization of diverse amyloidogenic immunoglobulin light chains revealed by hydrogen-deuterium exchange mass spectrometry

**DOI:** 10.64898/2026.01.07.698275

**Authors:** Daniele Peterle, Nicholas L. Yan, Elena S. Klimtchuk, Thomas E. Wales, Olga Gursky, Jeffery W. Kelly, John R. Engen, Gareth J. Morgan

**Affiliations:** Department of Chemistry & Chemical Biology, Northeastern University, Boston, MA, USA; Department of Chemistry, The Scripps Research Institute, La Jolla, CA, USA; Amyloidosis Center, Boston University Chobanian & Avedisian School of Medicine, Boston, MA, USA; Department of Pharmacology, Physiology & Biophysics, Boston University Chobanian & Avedisian School of Medicine, Boston, MA, USA; Section of Hematology and Medical Oncology, Department of Medicine, Boston University Chobanian & Avedisian School of Medicine, Boston, MA, USA

**Keywords:** Immunoglobulin light chain amyloidosis, AL amyloidosis, amyloid fibrils, Light chain kinetic stabilizers, drug design, protein stability and dynamics, HDX-MS, PLIMSTEX analysis

## Abstract

Immunoglobulin light chains, a component of antibodies, can misfold and aggregate to cause systemic AL amyloidosis. Aggregation, including amyloid fibril formation, requires unfolding of the full-length light chain from its native state, and in most cases aberrant proteolysis. Small molecules that bind to the native state of light chains to stabilize them against conformational excursions and proteolysis are under development as drug candidates for AL amyloidosis. Since each patient has a unique light chain sequence, a challenge for candidate stabilizer drugs is to bind multiple light chains and suppress their dynamics. Here, we used hydrogen-deuterium exchange measured by mass spectrometry to characterize the binding of six small molecule stabilizers to eleven different λ light chain proteins. Despite structural and dynamic differences among the light chains, the binding of the most efficacious stabilizer molecule led to increased protection from hydrogen exchange, consistent with reduced local and global unfolding. Protection upon binding was most prominent in residues within complementarity determining region 3 and framework region 4 of the light chain variable domains, which undergo major conformational changes enabling amyloid formation. Stabilizer binding also reduced the rate at which all light chains were cleaved by protease. These data show that these stabilizers suppress the range of conformational dynamics associated with light chain aggregation, supporting their therapeutic potential.

**Highlights:**

- Small-molecule kinetic stabilizers suppress conformational dynamics and aberrant proteolysis of diverse amyloidogenic full-length λ light chains.
- HDX-MS indicates conserved protection in regions forming the interface between the two variable domains of the light chain dimer, corresponding to residues that form the core of patient-derived amyloid fibrils.
- PLIMSTEX binding titrations show coupled binding, dimerization and stabilization at sub-micromolar kinetic stabilizer concentrations.
- Small molecule kinetic stabilization across different λ sequences supports kinetic stabilizers as a promising therapeutic strategy for AL amyloidosis.

## Introduction

Unfolding and subsequent aberrant proteolysis of full-length (FL) immunoglobulin light chain (LC) proteins are generally required for the formation of amyloid light chain (AL) deposits in the disease AL amyloidosis (Figure 1A), one of a spectrum of systemic amyloidoses (Buxbaum et al. 2024). Accumulation of amyloid fibrils derived from normally soluble LCs in multiple tissues leads to progressive and eventually fatal organ dysfunction (Sanchorawala 2024; Merlini et al. 2018). Soluble, misfolded LCs can also contribute to cardiotoxicity that is the major cause of mortality (Merlini et al. 2018). FL LCs are subunits of antibodies that normally function in the immune system to recognize antigens (Del Pozo-Yauner et al. 2023). Antibodies are covalent heterotetramers of two FL LCs and two heavy chains, which circulate in blood alongside “free” LC monomers and homodimers—FL LCs are produced in excess to enable efficient antibody secretion (Feige, Hendershot, and Buchner 2010). Overproduction of a single monoclonal free LC by an aberrant clonal cell population can cause LC fragment accumulation as amyloid fibrils within tissues. Each AL patient has an essentially unique LC sequence that forms amyloid (Morgan et al. 2025). The amino acid sequence of each LC appears to define its propensity to aggregate (Del Pozo-Yauner et al. 2023; Absmeier et al. 2023). Amyloid formation is also driven, in part, by increased concentration of the monoclonal free FL LC or fragments thereof, so existing therapies kill the clonal cells to reduce the total amyloidogenic LC concentration (Sanchorawala 2024; Merlini et al. 2018). Protein conformational control by small molecules that modulate protein assembly, folding and stability have become an important class of drugs, targeting proteins associated with multiple disease categories (Chiti and Kelly 2022). Kinetic stabilizers are a subset of these molecules, which selectively bind to the native state and slow the rate of protein unfolding. Kinetic stabilization of circulating LCs should protect residual free LCs from misfolding, aberrant proteolysis and aggregation, thus complementing cytotoxic chemotherapies (Morgan, Buxbaum, and Kelly 2021), and this concept is currently being tested in clinical trials.

**Figure 1:**
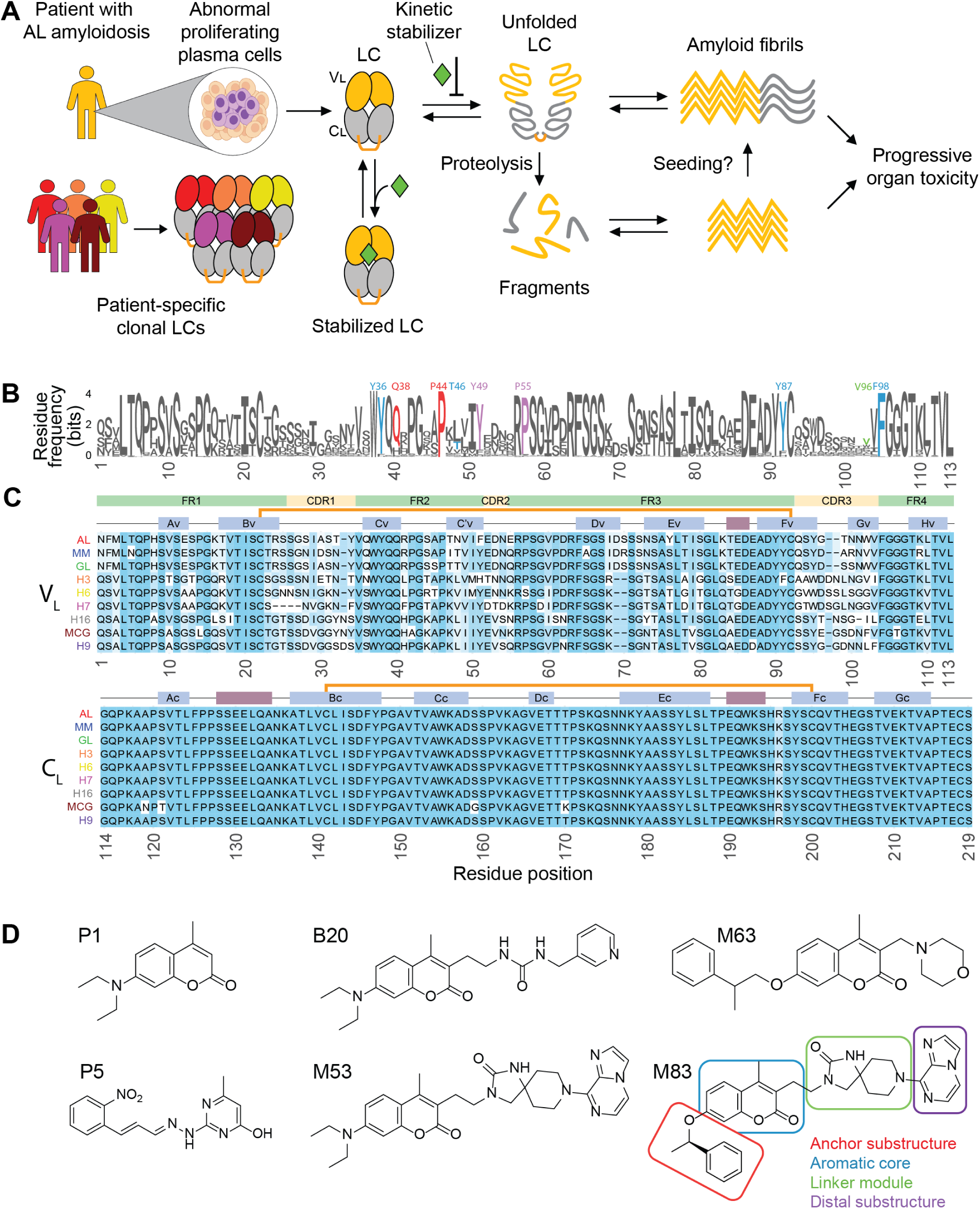
Kinetic stabilizers to inhibit aggregation of diverse antibody light chains. (A) Schematic representation of systemic AL amyloidosis pathogenesis and the mechanism of action of kinetic stabilizers. Unstable LCs are overexpressed by abnormal proliferating plasma cells and released into the bloodstream as homodimers, composed of a variable VL domain (in yellow) and a constant CL domain (in gray). An inter-molecular disulfide bond between cysteine residues at position 218 is observed in some LCs. Each clone produces a unique LC sequence. LCs can undergo unfolding, oligomerization, and proteolysis events, eventually leading to formation of fibrils that accumulate in various organs, most frequently the heart and kidney, resulting in progressive organ damage. Proteolysis can release aggregation-prone fragments that may act as templates or seeds for intact LCs. Kinetic stabilizers prevent fibril formation by stabilizing the dimeric native structure, thus reducing the concentration of aggregation-competent species. (B) Sequence logo plot of 554 VL domains of λ LCs taken from AL-Base. The height of each letter is proportional to its frequency in the alignment and the overall height of the column is proportional to the information content at that position. Amino acids interacting with kinetic stabilizers are indicated according to the numbering used in the original crystal structure (PDB code 6MG5) and color-coded according to the substructure of the M83 stabilizer to which they bind (see the structure of M83 in panel (D): red, anchor substructure; blue, aromatic core; green, linker module; purple, distal substructure). (C) Multiple sequence alignment of the 9 selected LCs explored in this study. The shading indicates the degree of conservation of amino acid residues. VL domain framework regions (FR1–4) and complementarity determining regions (CDR1–3) are shown in green and yellow, respectively. Blue and red boxes show the locations of the native secondary structure elements. Individual ß-strands are labeled. Intramolecular disulfide bonds are shown by orange lines. Note that the IMGT regions are defined to allow comparison between different immunoglobulin domains and do not correspond precisely to LC secondary structure elements. (D) Structures of the kinetic stabilizers used in this study. M83, the stabilizer molecule that showed the highest potency in LC stabilization, has its substructures highlighted.

There are two types of LC, κ and λ, of which λ LCs are more frequently involved in AL amyloidosis (Kourelis et al. 2017; Merlini et al. 2018; Morgan et al. 2025). FL LCs are secreted as 210–220-residue proteins that fold into two independent structural domains (Figure 1A), a variable (V_L_) domain and a constant (C_L_) domain (Del Pozo-Yauner et al. 2023; Feige, Hendershot, and Buchner 2010). Both domains have a ß-sandwich structure, termed an immunoglobulin (Ig) fold, stabilized by an intramolecular disulfide bond. The N-terminal variable domain (residues 1–113 in the alignment shown in Figures 1B and 1C) is involved in antigen binding when incorporated into antibodies via three highly polymorphic loops known as complementarity determining regions (CDR, Figure 1C). Other residue segments, defined as framework regions (FR), are more conserved. The C-terminal constant LC domain (residues 114-219) has a structural role within antibodies, and its sequence is highly conserved between LCs. Free FL LCs can form homodimers in circulation. Such free LC dimers have a parallel orientation where the V_L_ and C_L_ domains from one protomer interact with their counterpart in the other protomer (Figure 1A). Residues from ß-strands C_V_, C’_V_, F_V_ and G_V_ form a V_L_-V_L_ interface, while residues from ß-strands A_C_, B_C_, D_C_ and E_C_ form the C_L_-C_L_ interface. A cysteine residue near the C-terminus of C_L_ (referred to here as C218, relative to the alignment) can make an intermolecular disulfide bond within the homodimer. In patients, amyloidogenic FL LCs can form both monomers and homodimers (Connors et al. 2007; Andrich et al. 2017; Sternke-Hoffmann et al. 2020).

The structured cores of patient-derived amyloid fibrils primarily comprise residues from the V_L_ domain in non-native, cross-ß-sheet conformations (Radamaker et al. 2019). Isolated V_L_ domains can aggregate under mild conditions *in vitro*, whereas FL LCs remain soluble for extended periods under the same conditions and are much more resistant to aggregation (Morgan and Kelly 2016; Blancas-Mejía et al. 2015; Blancas-Mejía and Ramirez-Alvarado 2016). Amyloid-associated V_L_ domains tend to unfold and aggregate more readily than non-AL V_L_ domains (Wall et al. 1999; Hurle et al. 1994; Baden, Randles, et al. 2008; Kazman et al. 2020; Absmeier et al. 2023). Furthermore, amyloid-associated FL LCs are generally less conformationally stable than their non-AL counterparts and are more prone to local or global unfolding that enables aberrant endoproteolysis, releasing aggregation-prone fragments, including the V_L_ domain (Klimtchuk et al. 2010; Morgan and Kelly 2016; Oberti et al. 2017; Blancas-Mejía et al. 2017; Lavatelli et al. 2024). In some cases, proteolysis can occur after aggregation (Lavatelli et al. 2020). Amyloid fibrils formed by V_L_ domains can accelerate, or seed, the aggregation of full-length LCs (Blancas-Mejía and Ramirez-Alvarado 2016).

Collectively, these data suggest a role for the local conformational dynamics in sensitive protein regions. These regions have been proposed to involve: i) domain-domain interfaces in free LCs, such as the V_L_-V_L_ and V_L_-C_L_ interfaces (Rennella et al. 2019; Brumshtein et al. 2014; Baden, Owen, et al. 2008); ii) regions involving the internal disulfide bond in V_L_, a region that unfolds during amyloid formation (Radamaker et al. 2019; Klimtchuk et al. 2023); iii) amyloid-promoting regions (APRs), which are sequence regions with high intrinsic aggregation propensity that are proposed to initiate protein misfolding (Klimtchuk et al. 2023; Peterle et al. 2021; Goldschmidt et al. 2010); iv) mutation, misfolding or proteolysis of the C_L_ domains (Klimtchuk et al. 2010; Benson, Liepnieks, and Kluve-Beckerman 2015; Morgan, Usher, and Kelly 2017; Rottenaicher et al. 2023; Lavatelli et al. 2024). The roles of these sensitive regions in amyloid formation may differ for different LC sequences, complicating their therapeutic targeting.

Because LCs have diverse sequences and no natural small-molecule ligands, historically they epitomized a challenging drug target (Yan et al. 2023). Nontheless, we carried out a high-throughput screen that identified small molecules that protect folded FL LCs from undergoing conformational excursions enabling proteolysis, compounds that we call LC kinetic stabilizers; these molecules bind highly conserved residues at the V_L_-V_L_ interface of the LC homodimer (Morgan et al. 2019) (Figure 1B, colored residues). Subsequent structure-based medicinal chemistry efforts led to more potent kinetic stabilizers (Figure 1D) (Yan et al. 2020, 2021, 2022; Yan, Wilson, and Kelly 2023; Lederberg et al. 2024). Screening and optimization of hit molecules was mainly carried out using a single full-length LC protein, termed WIL-FL. A key challenge for ongoing drug development is to identify kinetic stabilizers that bind tightly and specifically to multiple LCs with diverse sequences. These kinetic stabilizers slow LC unfolding by strengthening the V_L_-V_L_ domain interface within the LC homodimer, which is expected to decelerate proteolysis and aggregation. However, the effects of small-molecule kinetic stabilizers on localized unfolding and proteolysis of a broader collection of FL LCs associated with amyloidosis have not been scrutinized experimentally.

To explore the conformational dynamics of multiple FL LCs we used hydrogen-deuterium exchange (HDX) mass spectrometry (MS). Previously, we used HDX-MS to investigate the native structure and dynamics of isolated V_L_ domains, FL LCs, and IgG proteins to identify specific regions that potentially contribute to protein misfolding enabling aberrant proteolysis (Peterle et al. 2021; Klimtchuk et al. 2023, 2024). In the current study we investigated localized unfolding in nine different FL LCs and the influence that the binding of small-molecule kinetic stabilizers have on these dynamics. The stabilizers explored were either screening hits (e.g. P1 and P5, Figure 1D) or structure-based designed stabilizers (e.g. B20, M63, M53 and M83. We provide molecular details about stabilizers’ mechanism of action and gain insights to guide the development of novel therapies for the treatment of AL amyloidosis.

## Materials and Methods

### Protein selection, expression and purification

Nine LC proteins selected for analysis are listed in Table 1. Amino acid sequences of these LCs are listed in the Supplementary Information. All residues are numbered according to the sequence alignment shown in Figure 1C. Domain boundaries and CDR residues are defined according to the IMunoGeneTics (IMGT) system (Lefranc et al. 2003). Cysteine 218 in the alignment, which forms an intermolecular disulphide bond, corresponds to residue 214 in the Kabat numbering system (Kabat et al. 1987) and residue 127 of the C-region in the IMGT numbering system. Variable and constant germline genes were assigned using IMGT’s tools for peptide and nucleotide analysis (Lefranc et al. 2018), as appropriate.

**Table 1:**
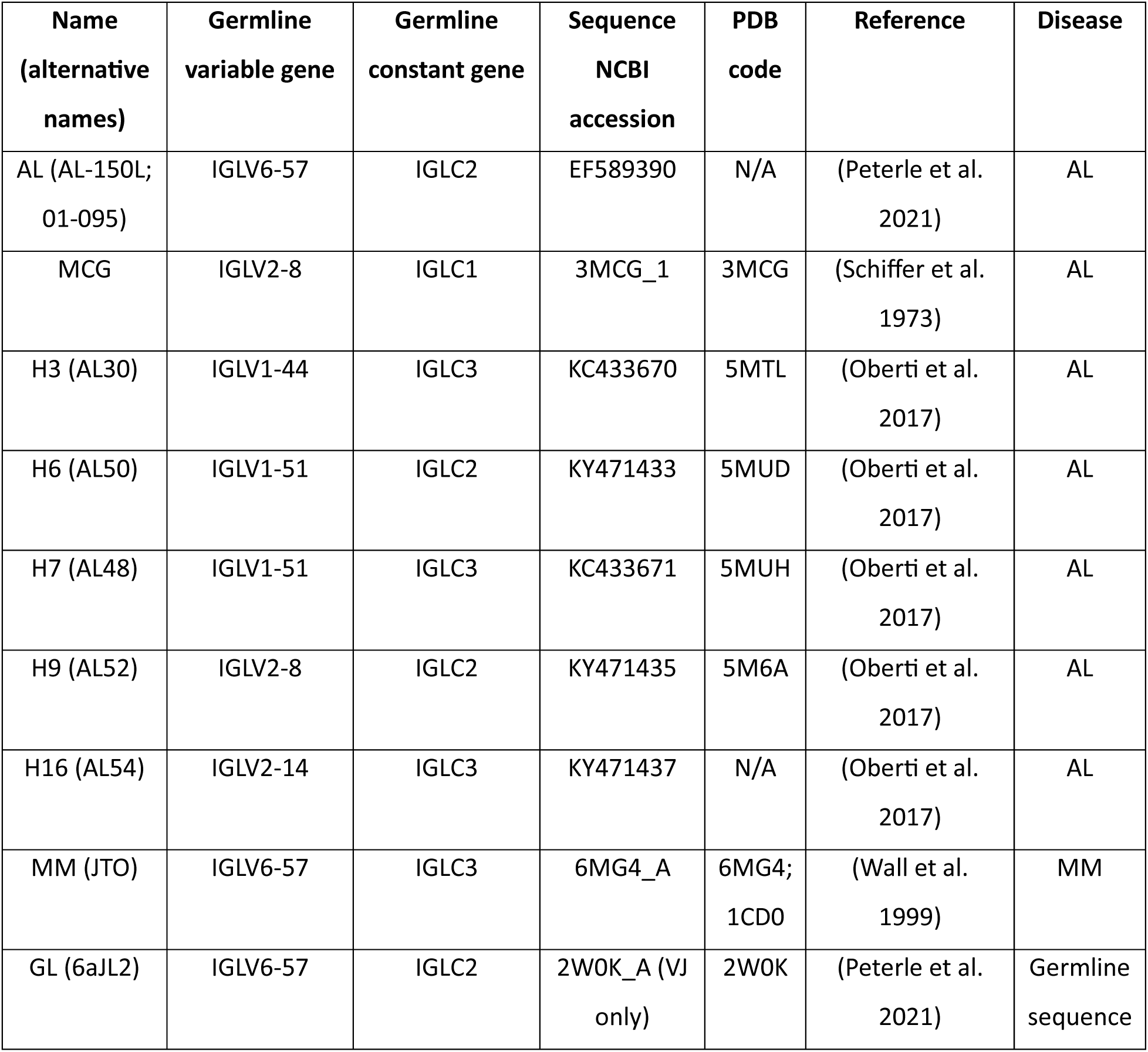
LC proteins used in the current study. Genes are defined according to the IMGT system. PDB identifiers are provided where experimental structures are available.

| Name<br>(alternative<br>names) | Germline<br>variable gene | Germline<br>constant gene | Sequence<br>NCBI<br>accession | PDB<br>code | Reference | Disease |
| --- | --- | --- | --- | --- | --- | --- |
| AL (AL-150L;<br>01-095) | IGLV6-57 | IGLC2 | EF589390 | N/A | (Peterle et al.<br>2021) | AL |
| MCG | IGLV2-8 | IGLC1 | 3MCG_1 | 3MCG | (Schiffer et al.<br>1973) | AL |
| H3 (AL30) | IGLV1-44 | IGLC3 | KC433670 | 5MTL | (Oberti et al.<br>2017) | AL |
| H6 (AL50) | IGLV1-51 | IGLC2 | KY471433 | 5MUD | (Oberti et al.<br>2017) | AL |
| H7 (AL48) | IGLV1-51 | IGLC3 | KC433671 | 5MUH | (Oberti et al.<br>2017) | AL |
| H9 (AL52) | IGLV2-8 | IGLC2 | KY471435 | 5M6A | (Oberti et al.<br>2017) | AL |
| H16 (AL54) | IGLV2-14 | IGLC3 | KY471437 | N/A | (Oberti et al.<br>2017) | AL |
| MM (JTO) | IGLV6-57 | IGLC3 | 6MG4_A | 6MG4;<br>1CD0 | (Wall et al.<br>1999) | MM |
| GL (6aJL2) | IGLV6-57 | IGLC2 | 2W0K_A (VJ<br>only) | 2W0K | (Peterle et al.<br>2021) | Germline<br>sequence |

Each LC listed in Table 1 was expressed as a full-length protein containing Cys218, which predominantly forms disulfide-linked dimers when expressed in *E. coli* (Peterle et al. 2021). We refer to these tabulated LCs as “AL LC”, “MCG LC”, etc. We also explored two additional proteins: the stand-alone V_L_ domain of the AL sequence (AL V_L_) and the AL C218S variant (Kabat: C214), where replacement of the cysteine prevents formation of the intermolecular disulfide bond. In total, 11 proteins were studied. Unless otherwise specified, experiments were carried out in phosphate buffered saline (PBS), defined here as 20 mM sodium phosphate buffer, pH 7.4, containing 150 mM NaCl. For proteins where no crystal structure was available, a model was generated using the AlphaFold 3 algorithm, implemented via www.alphafoldserver.com (Abramson et al. 2024). The relative orientations of the V_L_ domains within each model were similar but the “elbow” angle between the V_L_ and C_L_ domains varied; for clarity, only the V_L_ domains are shown when multiple structures are compared.

Variants of AL, MM and JTO LCs were expressed in *E. coli* with N-terminal polyhistidine tags and purified by affinity chromatography as previously described (Peterle et al. 2021). Untagged MCG, H3, H6, H7, H9 and H16 were expressed as inclusion bodies and purified by ion exchange and size exclusion chromatography, as previously described (Oberti et al. 2017; Yan, Wilson, and Kelly 2023). Protein purity was 95%+ verified by SDS-PAGE (Figure S1). Protein molar concentration is reported as monomer equivalents. For LCs characterized to date, stabilizers bind with a stoichiometry of one small molecule to two LC monomers.

### Small-molecule stabilizers

We tested six small-molecule kinetic stabilizers shown in Figure 1D and described in our previous work (Yan et al. 2023): P1, P5, B20, M53, M63 and M83. Numbering corresponds to that from the original paper (Yan et al. 2021). Letters indicate the origin of the molecules: P1 and P5 were originally identified as hits in a high-throughput screen (Morgan et al. 2019); B20, M53, M63 and M83 were designed and synthesized during medicinal chemistry efforts to improve stabilization efficacy (Yan et al. 2021, 2022). These molecules are elaborations on the coumarin scaffold of P1. B20 and M53 retain the original diethylamino substitution in position 7 of the coumarin heterocycle. M63 and M83 incorporate phenylethoxy modifications that were optimized for binding within the core of the LC dimer. Synthesis and purification of these molecules was carried out as previously described (Yan et al. 2021). Small molecules were solubilized in dimethyl sulfoxide (DMSO) vehicle and diluted to the desired concentration. DMSO concentrations were 1.5% during equilibration and 0.15% during deuterium labeling unless otherwise specified.

### Sequence analysis

All AL-associated λ LCs with complete V_L_ domain sequences were taken from AL-Base, available via https://wwwapp.bumc.bu.edu/BEDAC_ALBase/, and aligned according to the IMGT numbering system. Fully conserved gaps, which are used to facilitate structural comparisons between different types of immunoglobulin domains, were removed from the alignment, leaving 113 occupied positions. This alignment was visualized using the ggseqlogo (Wagih 2017) package in R v 4.2.2 (R Core Team 2022), and the binding site residues were highlighted manually.

The sequence-based consensus method AmylPred2 (Tsolis et al. 2013) was used to predict the amyloidogenic sequence propensities of the nine LCs under study. APRs are defined as segments predicted by 6 or more of the 10 algorithms available via the AmlyPred2 website.

### Hydrogen deuterium exchange mass spectrometry (HDX-MS)

At least two independent HDX-MS replicates were measured for each LC in accordance with current recommendations (Masson et al. 2019; Engen and Wales 2015). HDX-MS data have been deposited to the ProteomeXchange Consortium via the PRIDE repository (Perez-Riverol et al. 2019) with the dataset identifier PXD068745.

### Deuterium labeling

LCs were freshly diluted to 30 μM in biological replicates from stock concentrations in PBS, pH 7.4, prepared in H_2_O. Before labeling, the protein solutions were mixed 1-fold with a 300 μM solution of stabilizer molecule in PBS containing 3% (v/v) DMSO. Deuterium labeling commenced with a 10-fold dilution (18 μL) of the D_2_O labeling buffer (10 mM sodium phosphate, 150 mM NaCl, pD 7.4, 99.9% D_2_O) added to 2 μL of each 15 μM protein solution. After each labeling time (10 seconds, 1 minute, 10 minutes, 1 hour, 4 hours, 16 hours) at 20 °C, the labeling reaction was quenched with the addition of 20 μL of ice-cold quenching buffer [200 mM sodium phosphate, 4 M guanidine hydrochloride (GdnHCl), 0.72 M TCEP, pH 2.37, H_2_O] and analyzed immediately by liquid chromatography coupled mass spectrometry (LC-MS). The maximally deuterated samples of each protein that were used for back-exchange correction were prepared as described (Peterle, Wales, and Engen 2022). Protein solutions (15 μL at 15 μM) were lyophilized, resuspended in 7 M GdnHCl containing 50 mM DTT (15 μL), and heated at 90 °C for 5 min. After cooling to 20 °C, 18 μL of labeling buffer (as above) was added to 2 μL of cooled, denatured protein solution, and the exchange reaction was allowed to proceed at 50 °C for 10 min. These maximally deuterated protein samples were then cooled to 0 °C, exchange quenched with the addition of 20 μL of ice-cold quenching buffer (as above) and analyzed immediately by LC-MS.

### Mass spectrometry

LC-MS was performed with a Waters HDX system containing an HDX unit and two Acquity I-class UPLC pumps (Wales et al. 2008). Deuterated and control samples were digested online in the HDX cooling unit, where the digestion chamber was held at 15 °C, using an AffiPro Pepsin column (AffiPro, AP-PC-001, 2.1 mm × 20 mm). Peptides were trapped and desalted on a VanGuard PreColumn trap [2.1 mm × 5 mm, ACQUITY UPLC BEH C18, 1.7 μm, (Waters, 186002346)] for 3 minutes at 100 μL/min. Peptides were then eluted from the trap using a 5%–35% gradient of acetonitrile over 6 minutes at a flow rate of 100 μL/min, and separated using an ACQUITY UPLC HSS T3, 1.8 μm, 1.0 mm × 50 mm column (Waters, 186003535). The main cooling chamber of the Waters HDX unit, which housed all the chromatographic elements, was held at 0.0 ± 0.1 °C for the entire time of the measurements. The back pressure averaged ∼9000 psi at 0 °C and 5% acetonitrile 95% water, 0.1% formic acid. To eliminate peptide carryover, a wash solution (1.5 M GdnHCl, 0.8% formic acid and 4% acetonitrile) was injected over the pepsin column during each analytical run. Mass spectra were acquired using a Waters Synapt XS HDMSE mass spectrometer operated in ion mobility mode. The mass spectrometer was calibrated with direct infusion of a solution of Glu-fibrinopeptide (Sigma, F3261) at 200 femtomole/μL at a flow rate of 5 μL/min prior to data collection. A conventional electrospray source was used, and the instrument was scanned over the range 50 to 2000 m/z. The instrument configuration was: capillary voltage 2.5 kV, trap collision energy at 4 V, sampling cone at 35 V, source temperature of 80 °C, and desolvation temperature of 175 °C. All experiments were done under identical experimental conditions. In this experimental setup, the error in determining peptide deuterium levels was ±0.20 Da, as established using deuterated peptide standards.

### Data processing

Peptides were identified using PLGS 3.0.1 software (Waters, 720001408EN) with replicates of undeuterated samples used as controls. Raw MS data were imported into DynamX 3.0 (Waters, 720005145EN) and filtered with minimum consecutive products of 2 and minimum number of products per amino acid of 0.25. The peptides meeting the filtering criteria were further processed automatically by DynamX followed by manual inspection of all spectra and data processing. The amount of deuterium in each peptide was determined by subtracting the centroid mass of the undeuterated form of each peptide from that of the deuterated form, at each time point, for each condition. Correction for back exchange to report percent deuteration (%D) was done as described (Zhang and Smith 1993) according to the equation:

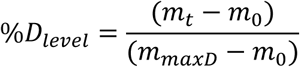

where m_t_ is the observed peptide centroid mass at a given labeling time-point t, m_0_ is the undeuterated peptide centroid mass, and m_maxD_ is the maximally deuterated peptide centroid mass.

Peptide-level “chiclet” plot heatmaps were used to visualize the HDX-MS data (Kerres et al. 2017). These visualizations directly show the acquired data, but because the analyzed peptides may not be equivalent between different proteins it can be difficult to make direct comparisons. To more easily visualize differences between LCs, we used the PyHDX web server to reduce the peptide deuteration level information to the single amino acid level (Smit et al. 2021). We assumed that both subunits of the LC dimers were equivalent. All measured and processed values can be found in the Excel supplemental data file. Residue-level percent deuteration values, corrected for back-exchange, were used to generate skyline plots, heat maps, difference heat maps and structural images. Data were plotted onto structural models of the appropriate LC using PyMOL (Schrödinger LLC). For a simple comparison between LCs, we calculated the area under the exchange curve (AUC) over the residues with measurable HDX data in all experiments, using a trapezoidal method.

### PLIMSTEX analysis of binding affinity

Binding affinity between small-molecule stabilizers and proteins was measured by HDX-MS using the PLIMSTEX approach (Zhu et al. 2003). Binding isotherms were generated by incubating 5 μM of protein (AL LC, C218S, or V_L_) with 2-fold serial dilutions of stabilizer molecules (500 to 1.9 μM) at 20°C in 10 mM sodium phosphate buffer, pH 7.4, containing 150 mM NaCl and 5% DMSO (v/v). After 30 minutes of equilibration at room temperature, 3 μL of the protein-ligand mixture was diluted with 57 μL of deuterium labeling buffer (10 mM sodium phosphate, 150 mM NaCl, pD 7.4, 99.9% D_2_O, final DMSO 0.25% v/v) and incubated for 1 hour and 30 minutes at 20°C. After dilution, protein concentration was 0.25 μM and small-molecule stabilizers ranged in concentration from 0.095 to 25 μM. The reaction was acid-quenched and immediately injected into the mass spectrometer for analysis. All experiments were performed in duplicate.

PLIMSTEX experiments were conducted on a Waters SELECT Series CyclicIMS system coupled to two I-Class UPLC pumps (Waters). The separation gradient was 5-35% acetonitrile over 3 minutes at a 100 μL/min flow rate. The mass spectrometer was calibrated using the Major Mix (Waters) and operated with the following parameters: capillary voltage 2.8 kV, trap collision energy 4 V, sampling cone 30 V, source temperature 80°C, and desolvation temperature 175°C.

For the PLIMSTEX data analysis of AL LC titration with M83, the peptide fragment ^101^NWVFGGGTKL^110^ (numbered according to the sequence alignment, equivalent to residues 98-107 of the AL LC sequence) was selected. This peptide was chosen because it contains residues that form the stabilizer binding site in WIL-FL, notably F104 (Morgan et al. 2019). Accordingly, it exhibited one of the strongest responses to kinetic stabilizer binding.

### Size exclusion chromatography

Protein (AL V_L_, 0.24 mg/mL, 20 μM) was incubated for 30 minutes at 37°C in the presence or absence of M83 (100 μM) in PBS, pH 7.4, containing 1 % (v/v) DMSO. Following incubation, the protein mixture was loaded onto a Superdex® 75 10/300 column (Cytiva) equilibrated with PBS, pH 7.4, and eluted using an AKTA Pure FPLC system (Cytiva) at a flow rate of 0.5 mL/min, while monitoring the absorbance at 280 nm. M83 was added at a 1:5 (mol:mol) ratio. The column was calibrated using protein standards (Sigma): blue dextran (2000 kDa), bovine serum albiumin (66 kDa), carbonic anhydrase (29 kDa), cytochrome C (12.4 kDa), and aprotinin (6.5 kDa).

### Limited proteolysis

LCs (0.2 mg/mL, ∼8.3 µM) were incubated in PBS, pH 7.4, containing 1% DMSO at 37°C with agitation at 450 rpm in the presence or absence of 100 µM of M83. Proteolysis was initiated by adding trypsin (Promega, 0.5 mg/mL stock concentration, ∼20.8 uM). At designated time intervals, aliquots (10 µg) of each proteolysis mixture were quenched with 0.2 % v/v formic acid and separated by denaturing electrophoresis on 10-20% acrylamide gradient Tris-Glycine gels under reducing conditions, followed by Coomassie staining. Intact LCs was quantified by densitometric analysis of the gel bands (Gel Analyzer v23.1, Istvan Lazar Jr. and Istvan Lazar Sr., available at www.gelanalyzer.com). Residual LC fractions were calculated for each time point. Proteolysis kinetics were measured over 24 h. Proteolysis rates were calculated by fitting the data to a single exponential decay, and areas under the curve (AUC) calculated using a trapezoidal method. These data were used to optimize a single timepoint experiment to determine the maximum protection afforded by kinetic stabilizer binding. Enzyme to substrate (E:S) ratios ranged from 1:16 to 1:64 (w:w) and incubation times from 5 hours to 24 hours, depending on the inherent proteolytic susceptibility of each LC variant.

## Results

### Structural dynamics vary among different full-length light chains

We investigated seven FL λ LC proteins whose sequences were originally identified in patients with AL amyloidosis, along with two FL λ LC control sequences (Figure 1, Table 1). These sequences are derived from germline genes which are over-represented in AL amyloidosis compared to the normal repertoire (Morgan et al. 2025; Kourelis et al. 2017). As in our prior study (Peterle et al. 2021), our controls were two non-amyloidogenic λ LCs: MM, originally identified in a patient with multiple myeloma but without amyloid deposition; and GL, the protein with the precursor germline sequence from which the AL and MM sequences are derived, IGLV6-57.

To identify APRs within the FL LC sequences we used the AmylPred2 server (Tsolis et al. 2013), which combines the results of multiple algorithms, producing a consensus score from 0 to 10. Similar regions of sequences were identified in each FL LC, although the strength of the predictions varied between sequences (Figure 2A). V_L_ domains harbored two (H6) to six (GL, MM, H16 and MCG) APRs (AmylPred2 score > 5). Two segments in the C_L_ domain, common to all nine FL LCs, were also predicted as APRs. These segments predominantly form ß-strands within the native LC structure (Figure 2A) but may drive self-association upon local or global unfolding.

**Figure 2:**
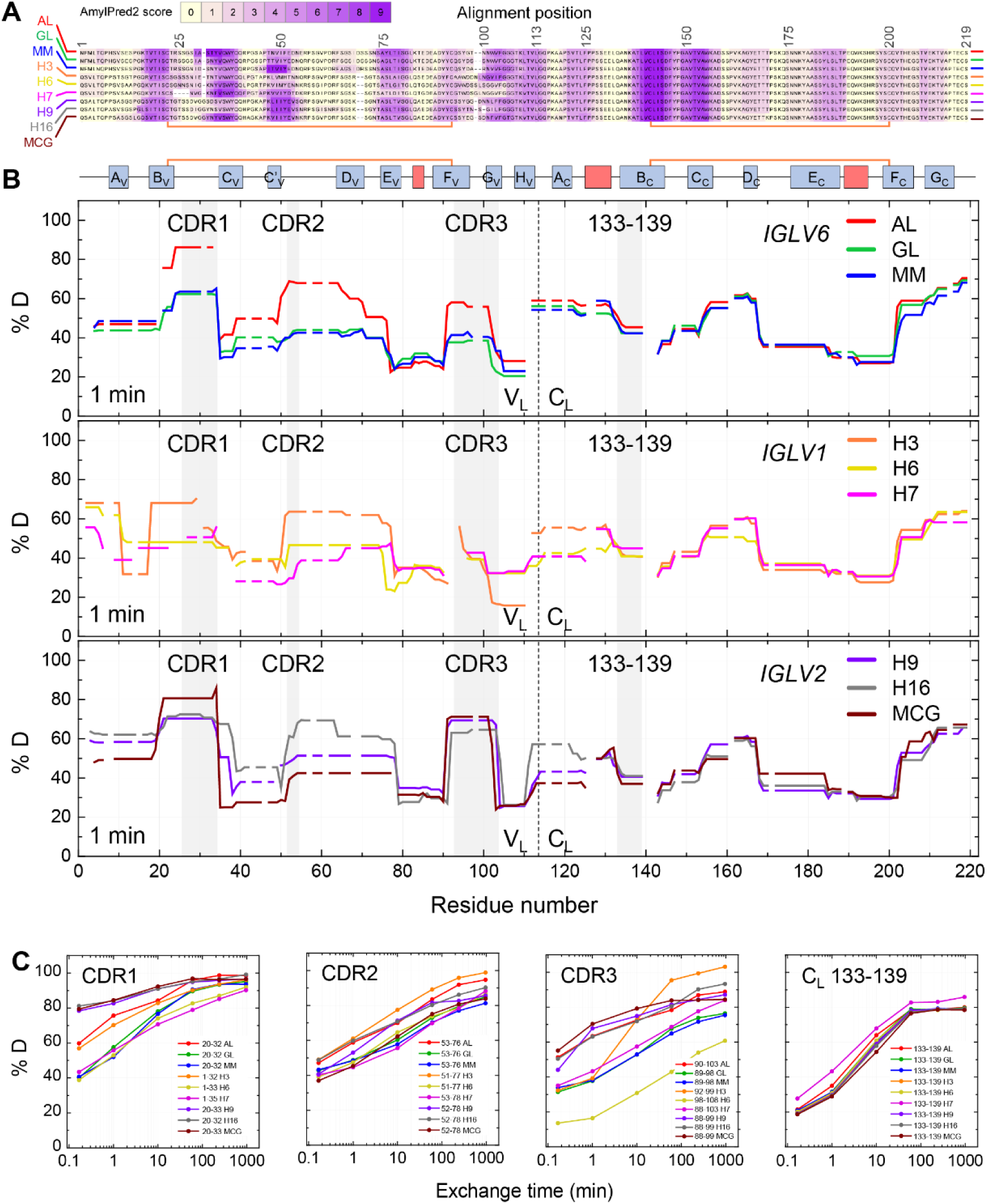
Hydrogen-deuterium exchange profiles of LCs. (A) Sequence alignment of the nine LCs under study, highlighting in purple the residues with amyloidogenic propensity as predicted by the AmylPred2 webserver. The shading represents the number of algorithms within AmylPred2 that identified each residue as amyloidogenic. (B) Skyline plot showing the percent of deuterium incorporation (%D) across the aligned LC amino acid sequences at 1-minute labeling time point. Gray shading represents CDRs 1, 2 and 3 within V_L_ and residues 133-139 within C_L_. Top, middle, and bottom panels correspond to LCs derived from variable gene families *IGLV6*, *IGLV1* and *IGLV2*, respectively. Dashed lines show the boundary between the variable and constant domains. The secondary structure scheme is shown at the top of the panels, with blue boxes representing ß-sheets and red indicating α-helices. Intramolecular disulfide bonds are shown by orange lines. (C) Deuterium uptake plots (%D) are shown for specific peptides covering regions of interest, including CDR1, CDR2, CDR3, and the constant region (C_L_). Each line corresponds to a specific LC variant, with peptide sequence numbers indicated. Note that these plots are not directly comparable due to sequence differences between the peptides. Complete HDX-MS results for all proteins and labeling times are reported in Figures S3, S6 and S7.

We first employed HDX-MS to investigate the local backbone conformational dynamics of FL LCs under identical conditions in the absence of kinetic stabilizer small molecules. HDX-MS provides detailed insights into solvent exposure and hydrogen bond stability by monitoring the exchange of backbone amide hydrogens with deuterons in the solvent (Konermann, Pan, and Liu 2011). The exchange was initiated under native-like conditions where the protein was folded. Greater deuterium incorporation reflected greater conformational dynamics and/or solvent exposure. Exchange was quenched at specific timepoints, from 10 sec to 16 h, by denaturation of the protein at acidic pH, where the average intrinsic rate of backbone amide HDX is minimal, followed by peptic digestion to generate peptides for LC-MS analysis. The extent of exchange at different time points was determined by measuring deuterium incorporation relative to that of the undeuterated protein, and was normalized relative to the experimental maximal deuterium uptake (maxD) observed for each peptide, as previously described (Peterle, Wales, and Engen 2022). For each LC, sequence coverage exceeded 95% (Figure S2).

To facilitate comparisons among different isoforms, HDX-MS results were visualized as skyline plots, which show percent deuteration at each residue interpolated from peptide-level exchange data. Figure 2B shows the data for 1 minute of exchange and Figure S3 shows other timepoints. No systemic differences between FL LCs with a histidine tag (AL, MM and GL) compared to other (untagged) FL LCs were observed by HDX (Figure 2B and S3). Figure 2C compares deuterium uptake kinetics for representative peptides in specific regions of interest, including CDR1, CDR2, CDR3, and the residue segment 133-139 from C_L_. The latter segment was selected because it partially overlaps with the amyloidogenic region 136-145 and encompasses a known proteolytic hotspot whose cleavage can generate a V_L_-containing LC fragment found in human amyloid deposits (Lavatelli et al. 2020; Mazzini et al. 2022; Lavatelli et al. 2024). We observed large variations in HDX within and between the FL LCs, mainly in the V_L_ domains, reflecting their variable sequences. As expected, the solvent-exposed CDR loops exhibit more deuteration, corresponding to the low structural protection of amides from HDX, as compared to the FRs, which contain more hydrogen-bonded β-sheets. The deuteration level in each region varied among different LCs. The least protected regions were adjacent to the APRs identified by AmylPred2. However, portions of these APRs showed low exchange indicating high protection by the native LC structure. This included high protection in C21 and C140 that are located in amyloidogenic segments and form internal disulfides in the V_L_ and C_L_ domains, respectively. The counterpart cysteine residues in the intermolecular disulfide bonds, C92 and C200, respectively, are located in regions of variable solvent exposure: C92 resides in a relatively exposed area near CDR3, whereas C200 is positioned within a more protected region.

As expected, the extent of C_L_ deuterium incorporation is similar for the nine LCs studied, reflecting the sequence and structural conservation of C_L_. Most differences in HDX were observed in the first 20 residues of C_L_ which are adjacent to the V_L_ domain. Within these residues, segments from AL, MM, GL, H3 and H16 showed lower structural protection (greater exchange) after 1 minute than those of H6, H7, H9 and MCG. These differences may be associated with differential interactions between the two domains.

Overall, no clear and strict correlation was identified between the observed deuteration levels in specific regions and the amyloid-forming properties of the LCs under study. However, compared to the non-amyloidogenic isoforms GL and MM, which share a very similar HDX profile, all amyloidogenic counterparts showed lower protection in the V_L_ domain, but not in the C_L_ domain. AL, H3, H9, and H16 exhibited generally rapid exchange across the entire V_L_, particularly in the CDRs. MCG showed greater deprotection in CDR1 and CDR3 at shorter time points, while FR3 exchanged to a greater extent at longer time points. H6 and H7 were more difficult to compare due to differences in peptide lengths and positions, but they exhibited increased uptake in FR4, FR1, and FR3 while remaining more protected in CDR1 and CDR2.

### Stabilizer binding reduces FL AL LC deuterium incorporation

We next investigated the effects of stabilizer molecules on deuterium uptake of AL LC. HDX-MS was carried out using 1.5 µM AL LC in the presence of 15 µM stabilizer. These concentrations are comparable with the circulating concentrations of free LCs (Katzmann et al. 2002) and an approved kinetic stabilizer drug for transthyretin amyloidosis (Cho et al. 2015). Figure 3A shows differences in normalized deuterium uptake of AL LC peptic peptides in the presence of P1, P5, B20, M53, M63 and M83, compared with the vehicle control (0.15% v/v DMSO), presented as peptide-level heatmaps (“chiclet plots”) across all labeling times. For clarity, only peptides from the V_L_ domain are shown; complete data are shown in Figure S4. Example data for the peptide covering residues 101–110 (^101^NWVFGGGTKL^110^, numbered according to the sequence alignment in Figure 1, corresponding to sequential positions 98–107 of AL LC) is shown in Figure 3B. This peptide comprises residues in CDR3 and FR4 and also forms part of the V_L_-V_L_ interface. No differences were detected at short labeling times; at longer times, increased stabilizer-induced protection from HDX was detected for four out of six stabilizer molecules (M83, B20, M53, P1). The region with the greatest change in protection from HDX was residues 92–110 (equivalent to sequential positions 89–107), which extends from C92 of the intra-domain disulfide bond through CDR3 to FR4. An additional site exhibiting less increased protection from HDX was observed between residues 34 and 48 (Figure 3A). Only minor stabilizer-induced changes were detected in C_L_ domain, which were most evident for residues 165-180 in the presence of M83 (Figure S4). Stabilizer molecules P5 and M63 did not alter deuterium uptake. The other four stabilizers resulted in increased protection from HDX in the order M83>B20>M53>P1. This is the same order as the small-molecules’ efficacy determined in the assay that was originally used for drug optimization, which measured protection of the WIL-FL LC from endoproteolysis (Morgan et al. 2019; Yan et al. 2021). Similar residues were protected from exchange by all four kinetic stabilizers.

**Figure 3:**
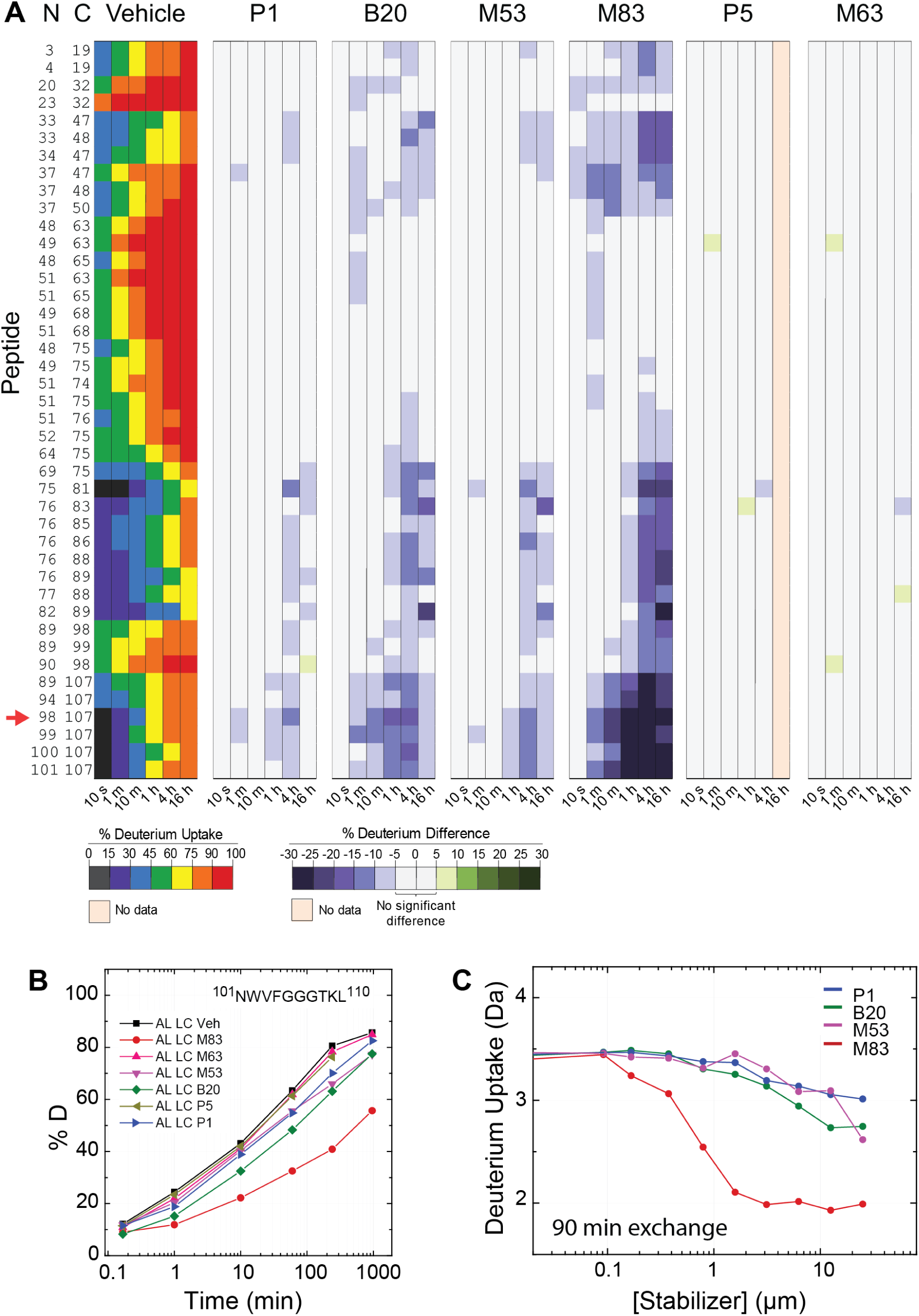
Differential stabilization of 1.5 µM AL LC by six small molecule stabilizers. (A) Peptide-level chiclet plot of deuterium uptake of FL AL LC in the presence or absence of 15 µM kinetic stabilizers. Only peptides originating from the V_L_ domain are shown; complete data are provided in Figure S4. “N” and “C” indicate the N- and C-termini, respectively, of the peptic peptides, numbered according to the sequence of AL LC. The AL LC peptide ^101^NWVFGGGTKL^110^ (corresponding to sequential positions 98-107) is highlighted with a red arrow. Vehicle (Veh) represents DMSO at 0.15% v/v during exchange. (B) Deuterium uptake curves for peptide ^101^NWVFGGGTKL^110^. (C) PLIMSTEX curves for peptide ^101^NWVFGGGTKL^110^ titrated with P1 (blue), B20 (green), M53 (magenta), and M83 (red) are shown as a function of the stabilizer molecule concentration.

To determine the relative efficacy of the small-molecule stabilizers, we measured deuterium uptake by AL LC (0.25 µM) as a function of stabilizer concentration. M83, B20, M53, and P1 were titrated from 0.2 to 25 µM. We analyzed the data using the PLIMSTEX approach (Protein Ligand Interaction by Mass Spectrometry, Titration, and EXchange) (Zhu et al. 2003; Zhu, Rempel, and Gross 2004). In a PLIMSTEX experiment the labeling time is fixed, and protein-ligand interaction is monitored through changes in deuterium incorporation across a titration series of the ligand. AL LC peptide ^101^NWVFGGGTKL^110^, described above, was selected for PLIMSTEX analysis. This peptide exhibits a large change in deuterium uptake upon stabilizer binding. The analysis was conducted at a labeling time of 1 hour and 30 minutes, chosen to capture the maximum observable protection from exchange while allowing the system to approach the near steady-state equilibrium of the exchangeable hydrogens (Zhu, Rempel, and Gross 2004). PLIMSTEX analysis of the HDX data indicated that M83 decreases deuterium uptake at concentrations 100-fold lower than that of the other stabilizer molecules (Figure 3C).

We chose to focus the remainder of our studies on M83, the most efficacious stabilizer, in order to investigate the mechanism by which M83 protects AL LC from exchange. We used two additional forms of the AL protein: AL LC C218S, which cannot form the intermolecular disulfide linking the C_L_ domains; and AL V_L_, the stand-alone V_L_ domain. Both these proteins equilibrate between monomer and non-covalent dimer in solution. At the concentrations used in HDX-MS, AL C218S is primarily dimeric and AL V_L_ is primarily monomeric (Peterle et al. 2021).

Figures 4A–4C show HDX-MS data from AL LC, AL C218S and AL V_L_, respectively, in the presence or absence of M83. Consistent with our previous studies, unliganded AL LC exhibited greater protection from HDX than AL C218S or AL V_L_ (difference heatmaps in the right-most columns of Figures 4B and 4C). These data are consistent with progressive stabilization associated with the presence of the C_L_ domain and intramolecular disulfide bond that has been observed in other FL LCs (Peterle et al. 2021; Klimtchuk et al. 2023; Rennella et al. 2019; Puri, Gadda, et al. 2025). Figure S5 shows the same data as peptide-level chiclet plots. Binding of the stabilizer molecule led to a large decrease in deuteration (i.e., an increase in HDX protection) observed from middle to long exchange times. For V_L_, this change in deuteration was no longer apparent at 16 hours, likely because both bound and unbound states of the protein had reached equilibrium with the solvent. Notably, this increased protection from HDX was observed in similar V_L_ segments of all three protein constructs. These segments were also more protected in AL LC than in AL C218S or AL V_L_, indicating that stabilizer binding mimics the effect of increased dimer formation that was previously observed for these proteins (Peterle et al. 2021). Unlike V_L_, the C_L_ domain showed only a small increase in protection upon stabilizer binding to FL LCs, mainly in residues 165–180, which are also more protected in AL LC than in AL C218S. The observed deuterium incorporation was only slightly reduced in M83-bound AL LC compared to M83-bound AL C218S. Accordingly, the change in deuterium incorporation upon M83 binding observed for AL C218S was greater than that observed for AL LC.

**Figure 4:**
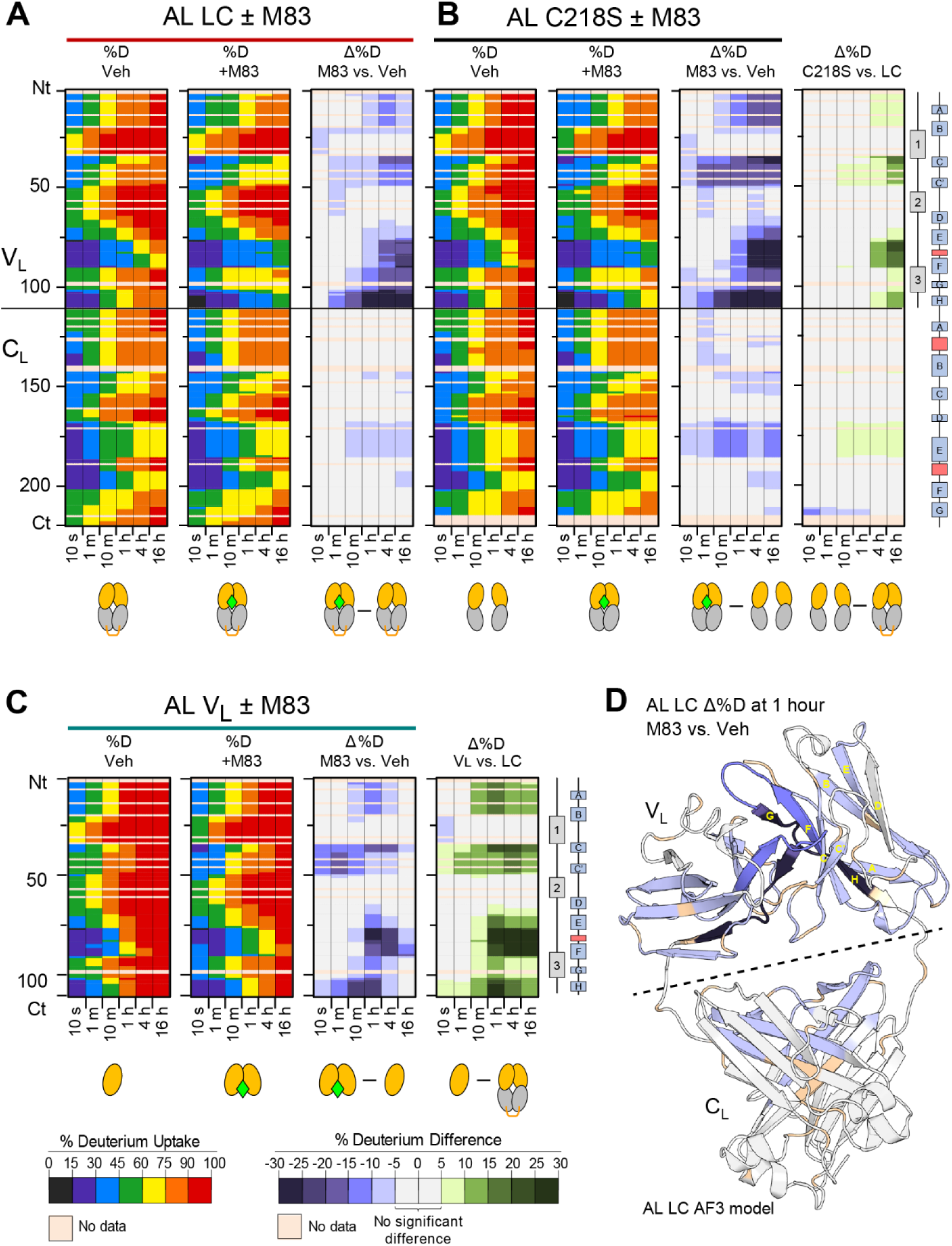
Effects of the kinetic stabilizer M83 binding to AL constructs monitored by HDX-MS. Residue-level heatmaps display percent deuterium incorporation (%D) for the proteins in the presence or absence of M83, as well as the corresponding differences in deuterium uptake (Δ%D). “Veh” represents DMSO vehicle at 0.15% v/v during exchange. Horizontal lines indicate the boundaries between the variable (V_L_) and constant (C_L_) domains. (A) AL LC. (B) AL C218S. (C) AL V_L_. Peptide-level data are provided in the Figure S5. Differences between constructs in the absence of M83 (C218S – LC and V_L_ – LC) are also shown. The protection induced by the kinetic stabilizer is greatest at peptides corresponding to the dimeric interface regions, but is not restricted to these residues. (D) Change in deuteration (Δ%D [(AL LC + M83) - AL LC]) after 1 hour of labeling mapped onto an AlphaFold 3 model of the AL LC dimeric structure. Regions of increased protection are shown in shades of blue. The names of the β-strands within the V_L_ domain are shown for one monomer.

We mapped the differences in AL LC deuteration in the presence and absence of M83 onto an AlphaFold3 model of AL LC, since no experimental structure of this FL LC is available (Figure 4D). Multiple backbone regions exhibited increased protection from HDX in the presence of M83. The greatest increase in protection was observed in peptides that form the V_L_-V_L_ interface. Altered exchange in this region is consistent with M83 binding in a similar site to that observed for other stabilizer molecules in co-crystal structures with LCs (Morgan et al. 2019; Yan et al. 2020, 2021; Yan, Wilson, and Kelly 2023; Lederberg et al. 2024). In addition, residues from the solvent-exposed ß-strands C_V_, C’_V_, F_V_ and H_V_, of the V_L_ domain also show increased protection upon M83 binding. C_L_ residues 165–180, which show a small increase in protection upon stabilizer binding, form the C_L_-C_L_ interface and are located close to the C_L_-V_L_ domain interface.

### Kinetic stabilizer binding enhances self-association of VL domains

To further investigate the interaction between stabilizer binding, dimerization and HDX, we applied the PLIMSTEX approach, as above, to measure the apparent affinity of M83 for the three AL constructs. We asked whether binding of M83 could force AL C218S and AL V_L_ to form a dimeric molecule with protection from HDX equivalent to that of the unbound or bound disulfide-linked dimer. M83 was titrated against each AL construct and the deuterium incorporation measured after 90 minutes. Raw mass spectra of peptide ^101^NWVFGGGTKL^110^ at varying concentrations of M83 are presented in Figure 5A, and the averaged centroid values of the spectra plotted against concentration of M83 are shown in Figure 5B. Without M83, deuterium incorporation for peptide ^101^NWVFGGGTKL^110^ differed significantly among the three AL constructs, reflecting differences in their intrinsic stabilities shown, consistent with previous experiments (Peterle et al. 2021). AL V_L_ exhibited the highest deuterium uptake (D_0_ = 6.34 Da, Figure 5A), AL C218S showed intermediate uptake (D_0_ = 5.20 Da), and AL LC showed the lowest uptake (D_0_ = 3.73 Da). M83 titration resulted in a concentration-dependent reduction in deuterium uptake across all three protein constructs, confirming specific binding. The maximum reduction in deuterium incorporation (ΔD, measured between the centroids of the m/z distributions) was observed at 25 µM M83, the highest stabilizer concentration investigated and 100-fold in excess of protein. The ΔD values for AL LC, AL C218S and AL V_L_ were -1.37 Da, -2.85 Da and 2.61, respectively. The change in deuteration was greater for the non-covalent constructs than for AL LC upon M83 binding. However, this quantitative analysis showed that neither AL C218S nor AL V_L_ reached the same protection from exchange as M83-bound AL LC. M83 binding was sufficient to reduce uptake of deuterium by AL C218S, but not AL V_L_, to below the level measured for unliganded AL LC. Fitting the observed ΔD to a unimolar binding model (1:1 LC dimer:ligand) yielded protection midpoints (EC_50_, the concentration of 50% maximal effect) of 0.65 μM for AL LC, 1.8 μM for AL C214S and 2.2 μM for AL V_L_ (Figure 5C; reported values are the mean of two independent measurements). These values are not directly comparable because the dimerization equilibria and exchange endpoints are different for the three AL constructs and hence, these values should not be treated as dissociation constants.

**Figure 5:**
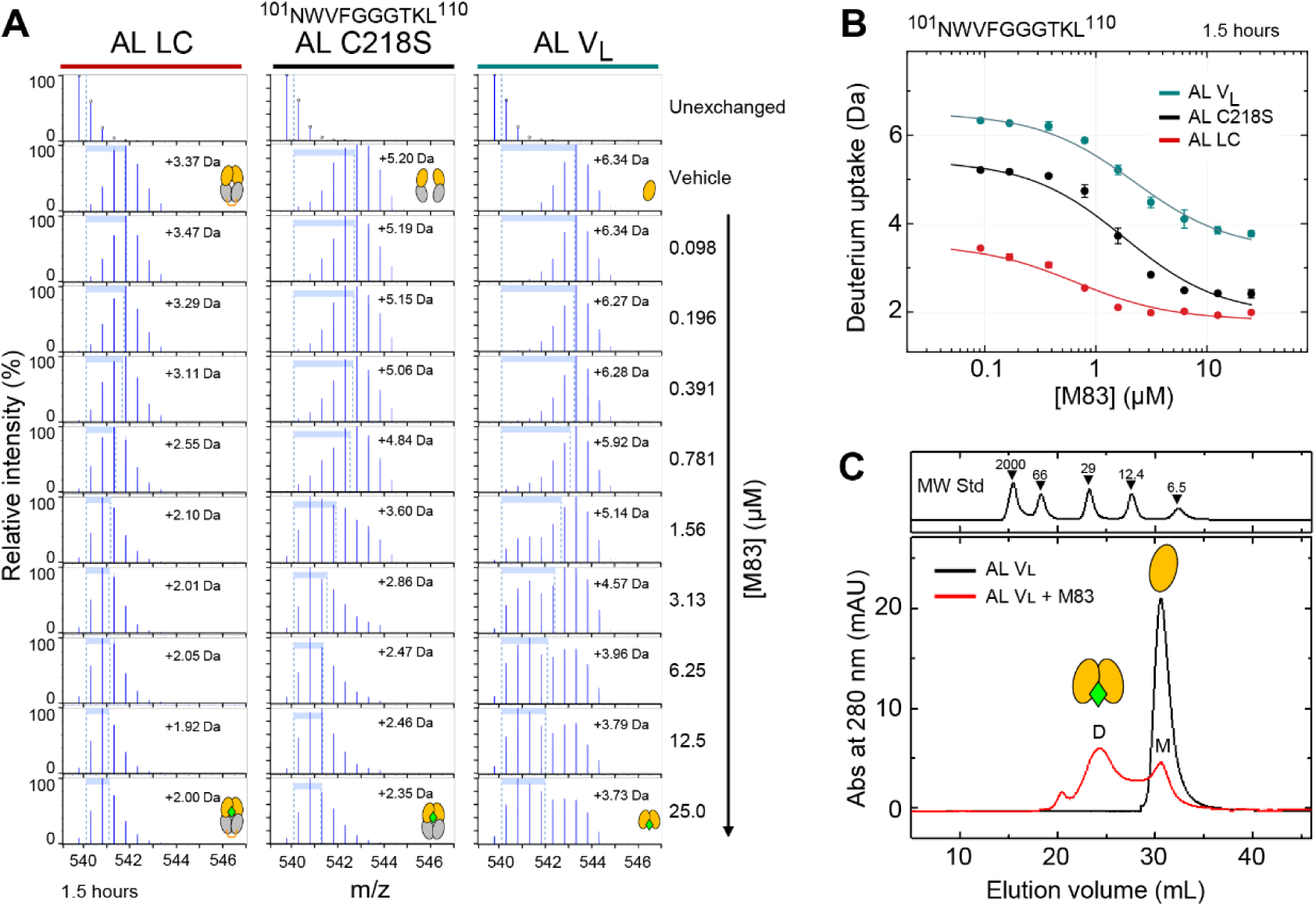
Stabilizer binding promotes LC dimerization. (A) Raw mass spectra of peptide ^101^NWVFGGGTKL^110^ from AL LC, AL C218S, and AL V_L_ in the presence of varying concentrations of M83 at a single time point of 90 minutes. Proteins concentration was 0.25 µM monomer equivalent. The blue peaks represent the isotopic distribution of the peptide, with centroid shifts from the undeuterated state indicating the average extent of deuterium incorporation (ΔD) (B) PLIMSTEX titration data for peptide ^101^NWVFGGGTKL^110^ from AL LC (red), AL C218S (black), and AL V_L_ (teal), showing measured deuterium uptake values (average of duplicates, where error bars show the range of the measurements) with corresponding fitted curves. (C) Size exclusion chromatography (SEC) of AL V_L_ in the absence (black) and presence (red) of 100 µM M83, illustrating its effect on dimer formation. M and D labels refer to the elution volume of the peaks corresponding to the monomeric and dimeric form of V_L_. Peaks corresponding to molecular weight standards are shown above.

The distributions of discrete m/z states observed in each experiment for peptide ^101^NWVFGGGTKL^110^ report on the ensembles of species within the exchange reaction, averaged over the 90-minute timecourse. AL LC spectra had similar distributions of peaks at all concentrations of M83, consistent with homogeneous exchange. The spectra of AL C218S and AL V_L_ were more complex, indicating the presence of multiple populations of exchanged species. Unliganded AL C218S had a broad distribution of states that becomes narrower as the concentration of M83 increases. AL V_L_ exhibited different behavior, where increasing M83 led to the appearance of a population with less incorporation of deuterium. We hypothesize that this behavior reflects ligand-induced dimerization of both non-covalent species. Under the maximum concentration of M83 tested, only a sub-population of AL V_L_ is stabilized.

We next asked whether kinetic stabilizer binding could enhance V_L_ domain dimerization. M83 increases the equilibrium population of V_L_ dimer, as determined by analytical size exclusion chromatography (Figure 5C). This observation is consistent with drug binding in the V_L_-V_L_ interfacial cavity, and with the previous observation that P1 binding favors dimerization of WIL-FL C218S (Morgan et al. 2019).

### Stabilizer molecules reduce hydrogen exchange and endoproteolysis of diverse LCs

We next asked whether binding of M83 alters the stability and dynamics of other FL LCs. We measured HDX kinetics for each FL LC (1.5 µM) in the presence and absence of 15 µM M83. Figure 6A shows the difference in deuteration between FL LCs with and without M83 as residue-level heatmaps, while Figure 6B shows the data mapped onto the dimeric V_L_ domains of structural models. All FL LCs exhibited reduced deuteration in the presence of M83, particularly between 1 to 4 hours of exchange (further data are provided in Figures S6 and S7). The protein segments showing the most significant stabilizer-induced protection from HDX were consistent across eight of the nine proteins, with the strongest effects observed in FR4 and, in some cases, CDR3. An exception was H6, which exhibited increased protection primarily within FR2. Only AL LC and GL LC exhibited increased protection within the C_L_ domain.

**Figure 6:**
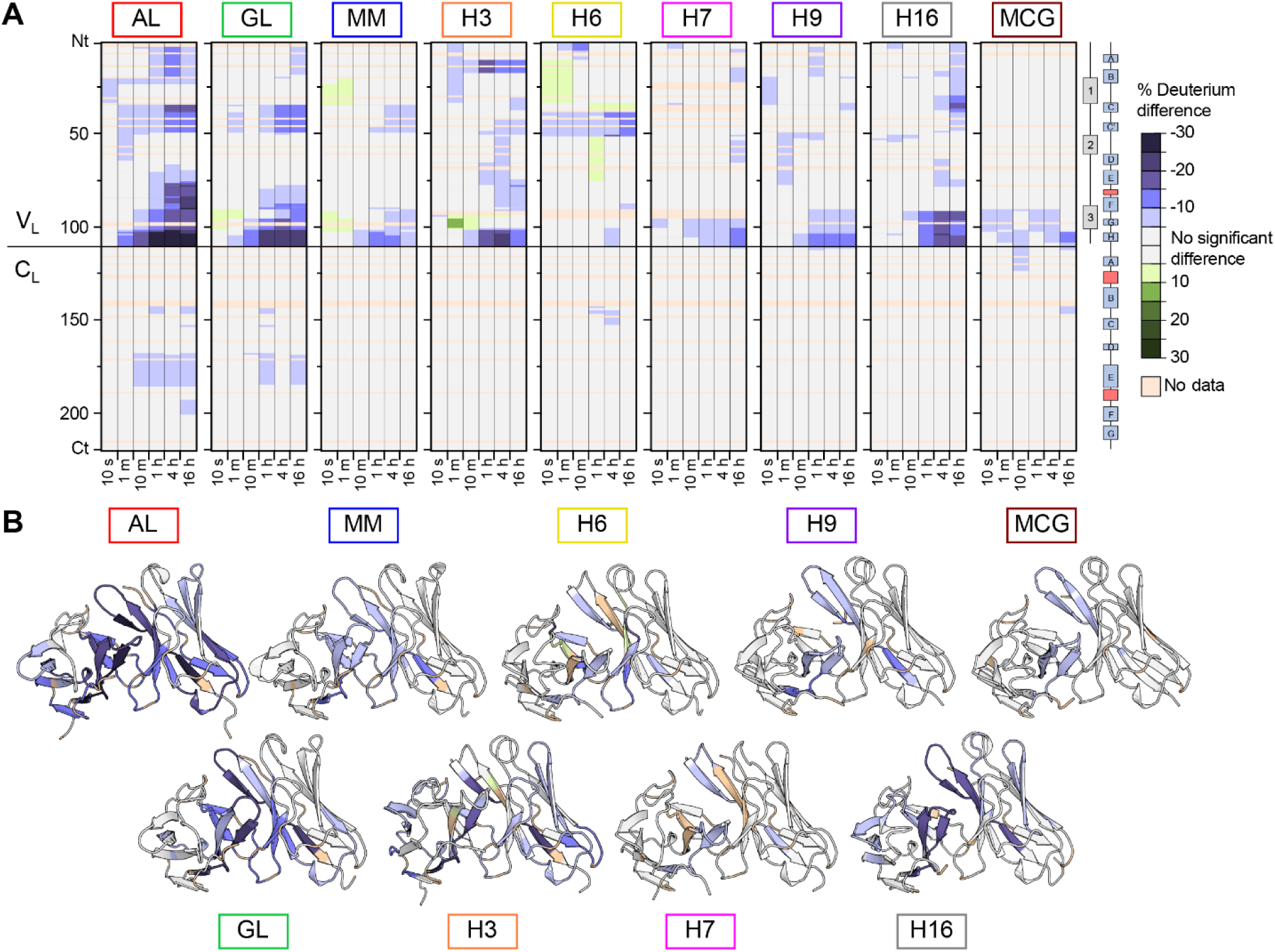
Effect of M83 binding on multiple LC proteins monitored by HDX-MS. (A) Single-residue aligned heatmaps of percent deuterium difference (Δ%D) across the nine LCs. The solid horizontal line indicates the boundary between the V_L_ and C_L_. Protection from deuterium exchange is observed for all LCs upon incubation with M83. Complete data are provided in Figure S6 and S7. (B) Structural mapping of Δ%D onto V_L_ dimer structures for each isoform highlights regions of stabilization upon M83 binding after 1 hour of exchange. The relative orientation of the V_L_ and C_L_ domains differs between the models, so for clarity only V_L_ is shown.

To further assess the structural impact of M83, we conducted limited proteolysis experiments using trypsin at neutral pH on FL LCs in the presence or absence of M83 (Figure 7). We initially measured timecourses for each FL LC under identical conditions, with timepoints ranging from 1 minute to 24 hours (Figure 7A and Figure S8). Although we were able to fit data for some FL LCs to exponential decay models (Figure 7A–B and Figure S8), this was not successful for all FL LCs. Therefore we calculated the area under the curve (AUC) for these data to define a simple metric for protease sensitivity (Figure 7A and Figure S8). For comparison, we calculated an equivalent AUC for HDX data (Figure 7B and Figure S8). Since the intrinsic proteolysis rates and the magnitude of M83-induced protection varied across the FL LCs, we tailored single timepoint proteolysis experiments by selecting optimized conditions for each FL LC (Figure 7D). The protease sensitivity of the FL LCs and the effect of M83 varied substantially. There was only a weak correlation between the intrinsic protease resistance of each FL LC and its protection from HDX (Pearson’s correlation coefficient *R* = 0.14, Figure 7E). Furthermore, the extent of protection from protease and HDX were weakly correlated, both when comparing the AUC for the proteolysis timecourses (*R* = -0.3, Figure 7F) and the optimized, single timepoint data (*R* = 0 069, Figure 7G). These observations indicate that limited trypsin proteolysis and HDX measure different aspects of stability, likely due to differential cleavage of unstructured peptide bonds within the folded states. However, both methods demonstrated stabilization of all FL LCs upon binding to M83.

**Figure 7:**
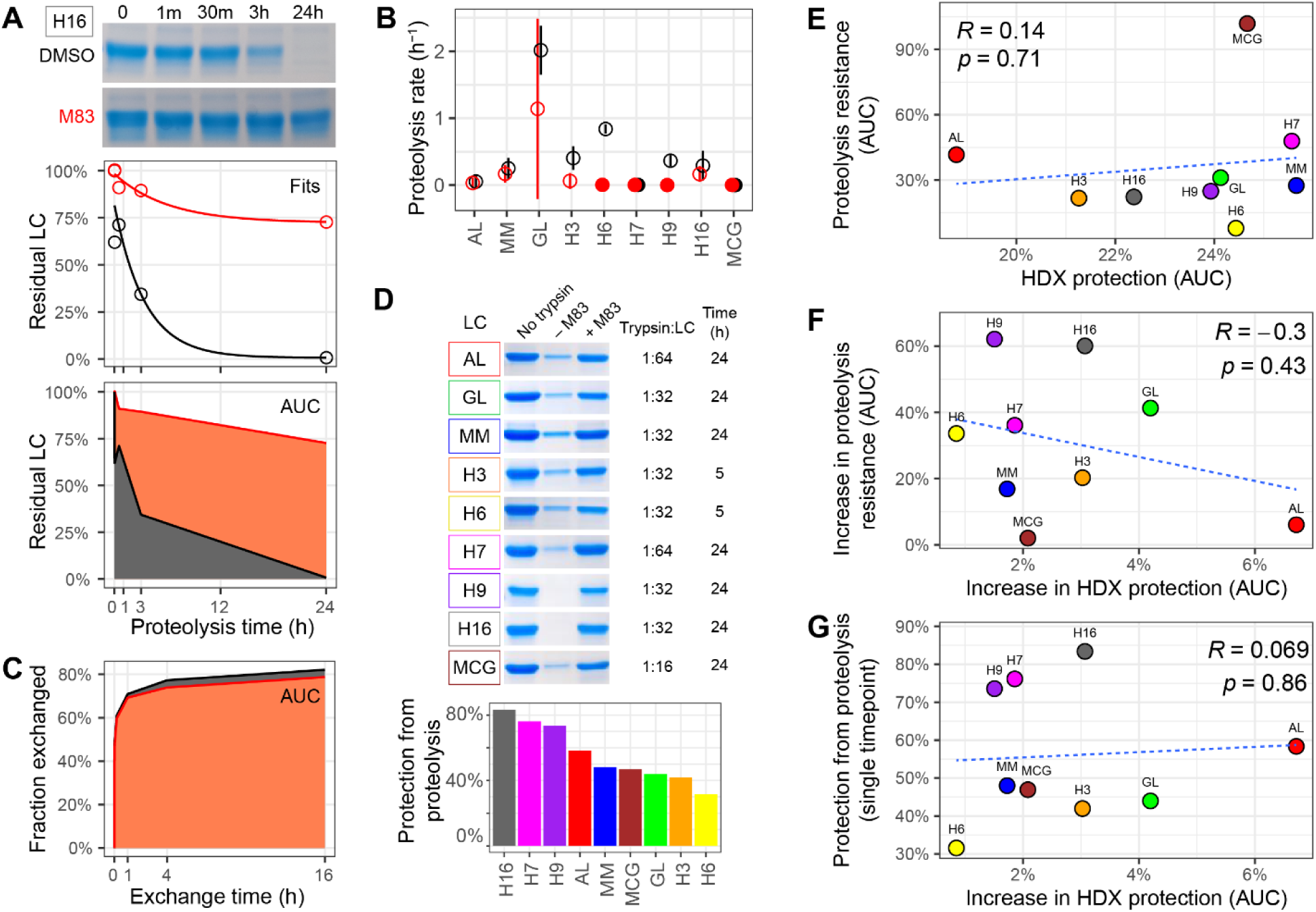
Effect of M83 binding on multiple LC proteins monitored by limited proteolysis. (A) Kinetics of digestion of H16 LC by trypsin in the presence (red) or absence (black) of M83, analyzed by SDS-PAGE. Data were analyzed by fitting to a single-exponential decay and by calculating the area under the curve (AUC). Data for other LCs are provided in Figure S8. (C) Proteolysis rates from fits of timecourse data to single exponential models. Hollow symbols and bars represent rates and fitting errors for successful fits (see Figure S8), solid symbols show data for which no rate could be determined. (B) AUC analysis of H16 HDX in the presence and absence of M83 for comparison with (A). (D) Single point proteolysis using optimized conditions to determine the maximal protection from protease afforded by M83 binding. Quantitation of the data is shown by the bars. (E–G) Correlations between proteolysis and HDX measurements of the intrinsic LC stability or stabilization imparted by M83. The blue dashed lines indicate linear regression lines with associated Pearson correlation coefficients (R) and significance values (p).

## Discussion

This work demonstrates the ability of small-molecule kinetic stabilizers to bind to multiple LCs and thereby suppress their dynamics and proteolytic susceptibility. LC dynamics, including local and global unfolding, enable self-association of non-native LCs to form amyloid, as well as proteolytic cleavage that yields more amyloidogenic V_L_-containing LC fragments. Binding of the most potent small molecule, M83, leads to a reduction in exchange of amide hydrogens with the solvent and decreased susceptibility to proteolytic cleavage for all nine LCs studied (Figures 6 and 7), despite their distinct sequences and dynamics (Figure 2). These data are consistent with the previous observation that small molecule binding stabilizes WIL-FL, the IGLV6-57-derived LC that was used for the high throughput screen and hit optimization to generate M83. The precursor germline genes represented by the LCs studied here, IGLV1-44, IGLV1-51, IGLV2-8, IGLV2-14 and IGLV6-57 together account for 386/847 (46%) of all amyloidogenic LCs and 386/621 (62%) of amyloidogenic λ LCs in the AL-Base repository (Morgan et al. 2025). By demonstrating that M83 stabilizes several LCs derived from the most commonly observed λ LC genes, our results support the hypothesis that pharmacologic stabilization of LCs is a promising therapeutic strategy that could complement existing and emerging approaches.

Consistent with our previous studies (Rennella et al. 2019; Peterle et al. 2021; Klimtchuk et al. 2023, 2024) and those of other groups (Puri, Gadda, et al. 2025; Weber et al. 2020; Rottenaicher et al. 2021; Puri, Palkar, et al. 2025), the extent of amide exchange varies across the sequence of the LCs. The V_L_ domains of the seven amyloidogenic proteins have distinct patterns of deuterium uptake. There are no clear patterns that could represent a “signature” of amyloid propensity other than our previous observations that AL LC is less protected from exchange than the non-amyloidogenic IGLV6-57-derived MM LC or GL LC. None of the LCs investigated here exhibited increased exchange in the C_V_ and C_V_’ strands (residues 38-49) that were identified as rapidly exchanging in the amyloidogenic IGLV6-57-derived LC AL55 which may be related to the altered V_L_–V_L_ interface observed for this specific LC (Puri, Gadda, et al. 2025). Instead, we observed more rapid HDX in the region encompassing CDR2 and strand D_V_ (Figure 2). In contrast, similar patterns of deuterium uptake were observed in all nine C_L_ domains, consistent with their similar sequences and structures. C_L_ domain mutations have been associated with instability and amyloid formation (Benson, Liepnieks, and Kluve-Beckerman 2015; Rottenaicher et al. 2023; Rennella et al. 2019), but our data suggest that in the absence of such mutations, the dynamics of the unmutated C_L_ sequences are indistinguishable within free LC dimers. Overall, these results and our previous study of κ LCs (Klimtchuk et al. 2023) are consistent with the hypothesis that multiple types of sequence and structural changes can lead to amyloidogenic LCs.

We asked whether stabilizer binding would lead to similar structural changes in LCs with different local dynamics. The stabilization imparted to all tested LCs is consistent with similar modes of binding between LCs, as observed in co-crystal structures of stabilizers with MM and H9 (Yan et al. 2021; Lederberg et al. 2024; Yan, Wilson, and Kelly 2023). However, our analysis only measures the effect of stabilizer binding on HDX and cannot directly report on the conformation of the LC or small molecule. It is possible that M83 can bind in different conformations to that observed for other stabilizers and LCs, particularly in the case of H6, which has a different pattern of differentially-protected regions than the other LCs (Figure 6).

We hypothesize that stabilizer binding enhances structural coupling between the constituent domains of a full-length LC dimer (Rennella et al. 2019), so that both global and local structural fluctuations are reduced. In agreement with this idea, stabilizer binding enhances dimerization and increases protection in both the residues that define the binding pocket and residues forming the dimeric interface, such as strand A, FR2, strands E and F, CDR3 and J-region of the V_L_ domain as well as strand E of the C_L_ domain (Figures 4–6). On the other hand, relatively little change in protection was observed for the most labile protons in the loops of CDR1 and CDR2, indicating that these residues do not form new hydrogen bonds. However, the global protection from protease digestion afforded by stabilizer binding indicates that these mobile regions are not susceptible to trypsin proteolysis without local or global unfolding.

Importantly, M83 binding increased local protection in CDR3 and FR4 regions in all proteins but H6 (Fig. 6A). CDR3 is adjacent to C92 that forms an internal disulfide (C21–C92) in V_L_. The structure around this conserved disulfide must fully unfold to enable the relative backbone rotation by 180 degrees, from parallel in native V_L_ to antiparallel in amyloid (Radamaker et al. 2019). This backbone rotation likely provides a major contribution to the kinetic barrier for LC misfolding in amyloid; if so, suppression of local backbone dynamics in the regions adjacent to the C21–C92 disulfide, particularly the highly dynamic CDR1 and CDR3, is expected to decelerate amyloid formation (Klimtchuk et al. 2023). Therefore, M83-induced stabilization of CDR3 and adjacent regions in most LCs explored suggests M83 should suppress LC amyloid formation.

The largest reduction in deuterium uptake induced by M83 was observed for AL LC (Figure 6A). Other amyloidogenic LCs showed a reduced level of stabilization, which may be due to the optimization of stabilizers for binding to IGLV6-57-derived LCs. The reduced protection afforded to GL LC and MM LC by stabilizer binding appears to be due to their increased stability in the absence of the small molecule. Binding of M83 is not sufficient to globally reduce the level of exchange of AL LC to that of the non-amyloidogenic LCs (Figure 6 and Figures S5–S6). However, the stabilization of AL LC’s structured regions by M83 reduced their exchange to a level similar to that observed for equivalent regions of unbound GL LC. The M83-bound states of AL LC and AL C218S exhibited similar exchange kinetics (Figure 4A and 5), where the M83-bound state of AL C218S was slightly less stable than M83-bound AL LC but more stable than unbound AL LC. We therefore suggest that stabilizer binding can compensate for the loss of stability associated with reduction of the intermolecular disulfide bond, and that the effect of stabilizers will not be limited to disulfide-linked dimers in patients.

The major limitation of our study is that these nine LCs represent only a fraction of the vast potential diversity of light chains. Although we expect that the stabilizer can bind to most λ LCs, it is not clear how much stabilization will be imparted in each case and whether this will be sufficient for clinical efficacy. Coumarin-based stabilizers including M83 bind relatively weakly to κ LCs (Morgan et al. 2019; Yan et al. 2021), which account for approximately 20% of AL amyloidosis clones (Morgan et al. 2025). Furthermore, it is not clear how the dynamics that are measured by HDX relate to amyloid propensity, since there is no clear correlation between the patterns of exchange protection observed for each LC and its pathogenicity. Our data will support further work to identify such mechanistic details.

Stabilization of LCs is a novel and potentially valuable therapeutic approach that could complement existing therapies. This work demonstrates that kinetic stabilizers can bind to multiple LCs and reduce the conformational dynamics associated with proteolysis and aggregation, which is essential for their broader utility. In combination with our previous work developing stabilizers with increased affinity (Yan et al. 2021) and specificity for LCs in blood (Lederberg et al. 2024), the current study supports further development of these molecules as therapeutic candidates.

## Declaration of interests

NLY, JWK and GJM are authors on a patent that describes the use of kinetic stabilizers as a potential therapy for AL amyloidosis. JWK is a founder and board member of Protego Biopharma.

## Abbreviations

AL: amyloid light chain
CDR: complementarity determining region
C_L_: light chain constant domain
FL: full-length
FR: framework region
HDX: hydrogen-deuterium exchange
GL: germline
LC: immunoglobulin light chain
MM: multiple myeloma
MS: mass spectrometry
SDS PAGE: sodium dodecyl sulfate polyacrylamide gel electrophoresis
ThT: thioflavin T
V_L_: light chain variable domain.

## Acknowledgements

This work was supported by National Institute of Health Awards R01-GM067260, R01-GM135158 and R01-HL157566; the Wildflower Foundation; and the Boston University Amyloid Research Fund.

## Supplementary Appendix

### Protein sequences

>AL

NFMLTQPHSVSESPGKTVTISCTRSSGSIASTYVQWYQQRPGSAPTNVIFEDNERPSGVPDRFSGSIDSSSNSAYLTISGLKTEDEADYYCQSYGTNNWVFGGGTKLTVL

GQPKAAPSVTLFPPSSEELQANKATLVCLISDFYPGAVTVAWKADSSPVKAGVETTTPSKQSNNKYAASSYLSLTPEQWKSHRSYSCQVTHEGSTVEKTVAPTECS

>GL

NFMLTQPHSVSESPGKTVTISCTRSSGSIASNYVQWYQQRPGSSPTTVIYEDNQRPSGVPDRFSGSIDSSSNSASLTISGLKTEDEADYYCQSYDSSNWVFGGGTKLTVL

GQPKAAPSVTLFPPSSEELQANKATLVCLISDFYPGAVTVAWKADSSPVKAGVETTTPSKQSNNKYAASSYLSLTPEQWKSHRSYSCQVTHEGSTVEKTVAPTECS

>MM

NFMLNQPHSVSESPGKTVTISCTRSSGNIDSNYVQWYQQRPGSAPITVIYEDNQRPSGVPDRFAGSIDRSSNSASLTISGLKTEDEADYYCQSYDARNVVFGGGTRLTVL

GQPKAAPSVTLFPPSSEELQANKATLVCLISDFYPGAVTVAWKADSSPVKAGVETTTPSKQSNNKYAASSYLSLTPEQWKSHKSYSCQVTHEGSTVEKTVAPTECS

>H3

QSVLTQPPSTSGTPGQRVTISCSGSSSNIETNTVNWYQQLPGTAPKLVMHTNNQRPSGVPDRFSGSRSGTSASLAIGGLQSEDEADYFCAAWDDNLNGVIFGGGTKLTVL

GQPKAAPSVTLFPPSSEELQANKATLVCLISDFYPGAVTVAWKADSSPVKAGVETTTPSKQSNNKYAASSYLSLTPEQWKSHKSYSCQVTHEGSTVEKTVAPTECS

>H6

QSVLTQPPSVSAAPGQKVTISCSGNNSNIGKNYVSWYQQLPGRTPKVIMYENNKRSSGIPDRFSGSKSGTSATLGITGLQTGDEADYYCGVWDSSLSGGVFGGGTKVTVL

GQPKAAPSVTLFPPSSEELQANKATLVCLISDFYPGAVTVAWKADSSPVKAGVETTTPSKQSNNKYAASSYLSLTPEQWKSHRSYSCQVTHEGSTVEKTVAPTECS

>H7

QSVLTQPPSVSAAPGQKVTISCSNVGKNFVSWYQQFPGTAPKVVIYDTDKRPSDIPDRFSGSKSGTSATLDITGLQTGDEADYYCGTWDSGLNGGVFGGGTKVTVL

GQPKAAPSVTLFPPSSEELQANKATLVCLISDFYPGAVTVAWKADSSPVKAGVETTTPSKQSNNKYAASSYLSLTPEQWKSHKSYSCQVTHEGSTVEKTVAPTECS

>H9

QSALTQPPSASGSPGQSVTISCTGTSSDVGGSDSVSWYQQHPGKAPKLIIYEVSQRPSGVPNRFSGSKSGNTASLTVSGLQAEDDADYYCSSYGGDNNLFFGGGTKVTVL

GQPKAAPSVTLFPPSSEELQANKATLVCLISDFYPGAVTVAWKADSSPVKAGVETTTPSKQSNNKYAASSYLSLTPEQWKSHRSYSCQVTHEGSTVEKTVAPTECS

>H16

QSALTQPASVSGSPGLSITISCTGTSSDIGGYNSVSWYQQHPGKAPKLIIYEVSNRPSGISNRFSGSKSGYTASLTISGLQAEDEADYYCSSYTNSGILFGGGTELTVL

GQPKAAPSVTLFPPSSEELQANKATLVCLISDFYPGAVTVAWKADSSPVKAGVETTTPSKQSNNKYAASSYLSLTPEQWKSHKSYSCQVTHEGSTVEKTVAPTECS

>MCG

QSALTQPPSASGSLGQSVTISCTGTSSDVGGYNYVSWYQQHAGKAPKVIIYEVNKRPSGVPDRFSGSKSGNTASLTVSGLQAEDEADYYCSSYEGSDNFVFGTGTKVTVL

GQPKANPTVTLFPPSSEELQANKATLVCLISDFYPGAVTVAWKADGSPVKAGVETTKPSKQSNNKYAASSYLSLTPEQWKSHRSYSCQVTHEGSTVEKTVAPTECS

### Supplementary Figures

**Figure S1.**
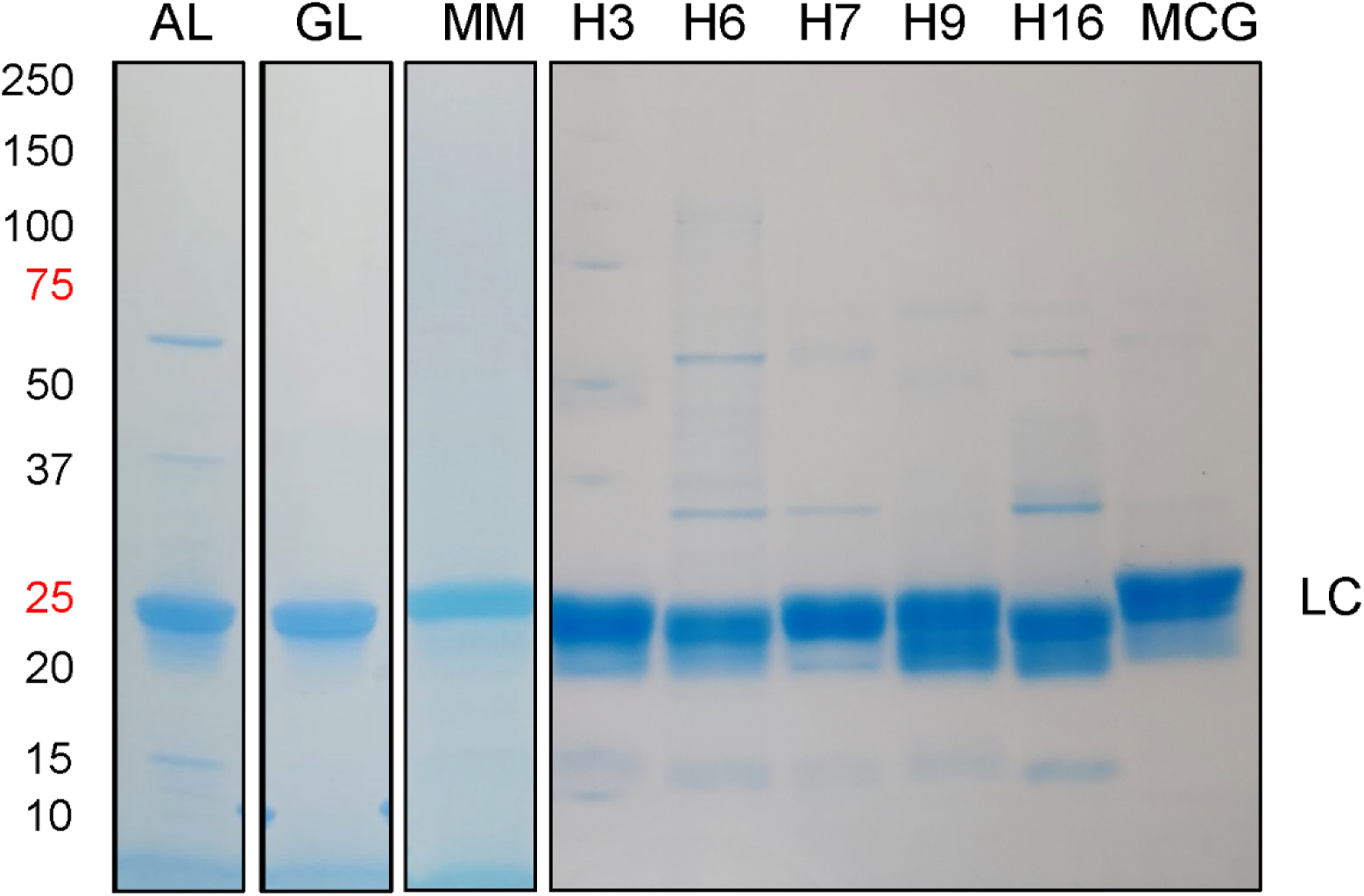
SDS-PAGE characterization of the nine full-length LCs used in this study. Proteins were run on a 10–20% acrylamide gel under reducing conditions and the gels were stained with Coomassie. 7 μg of each protein sample was loaded per lane. Molecular weight markers are indicated on the left. The secondary bands underneath the main LC band appear to be alternative conformers the main LC protein, since no truncated species were observed by mass spectrometry.

**Figure S2.**
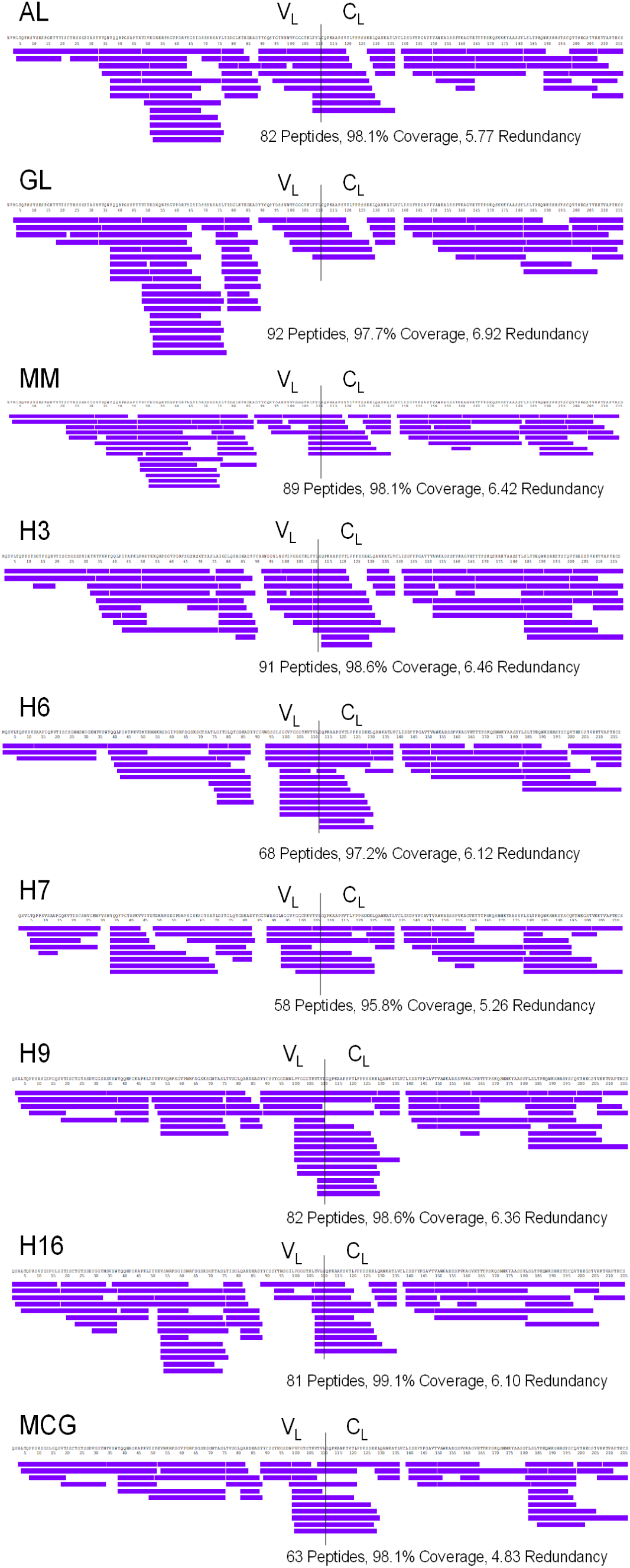
Sequence coverage maps of LCs constructs used in this study. Horizontal purple bars below LC sequence indicate the peptides for which HDX was followed. Vertical lines mark the separation between the V_L_ and the C_L_. The number of peptides, sequence coverage (%), and average redundancy values are reported for each LC.

**Figure S3.**
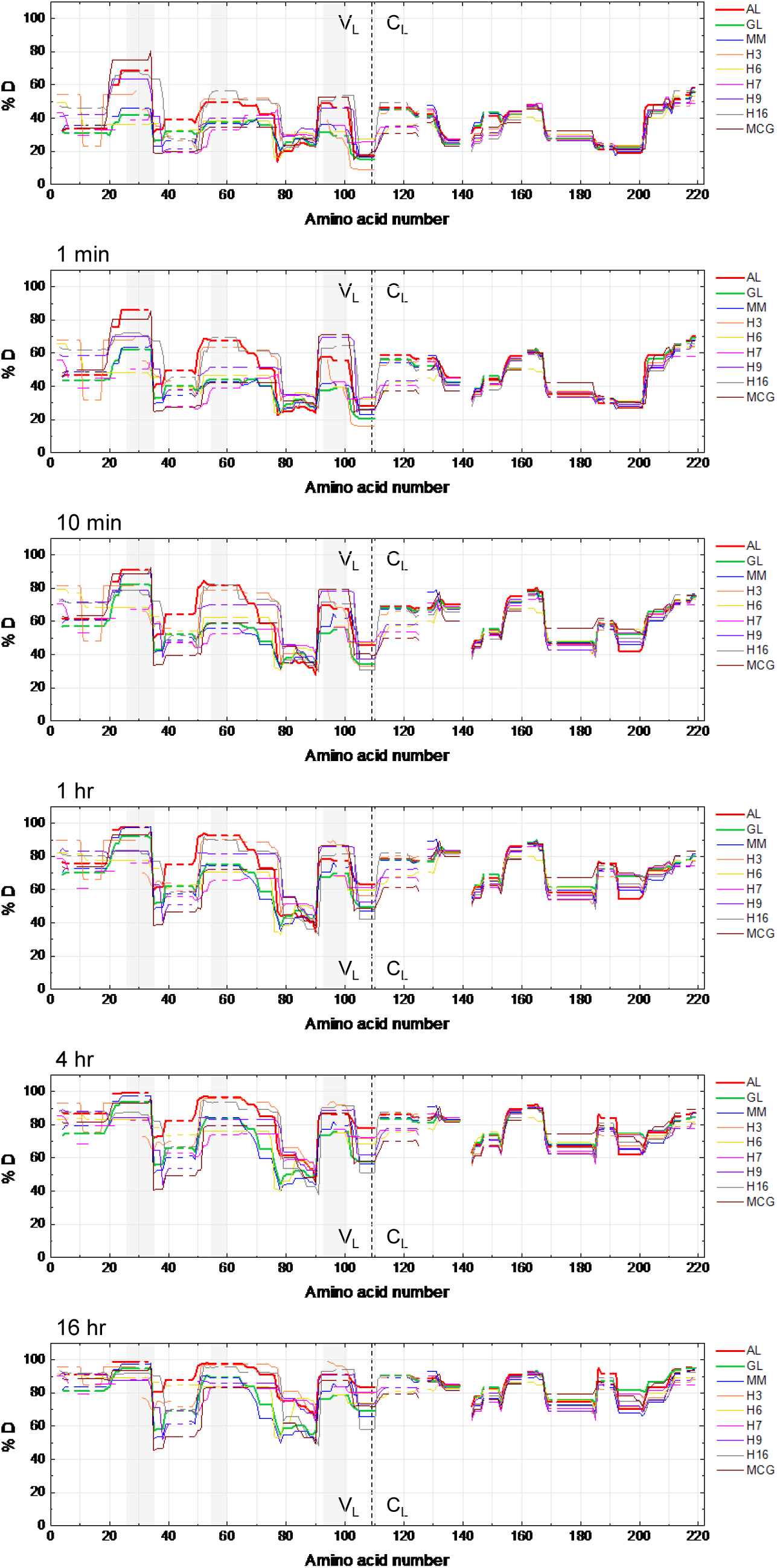
Single-residue aligned skyline plots for all light chains (LCs) at six different labeling time points: 10 seconds, 1 minute, 10 minutes, 1 hour, 4 hours, and 16 hours. Percent deuterium incorporation (%D) is plotted against amino acid number, with vertical dashed lines indicating the boundary between V_L_ and C_L_ domains. Gray boxes indicate the CDR regions. Each line represents a specific LC isoform, defined in the legend on the right.

**Figure S4.**
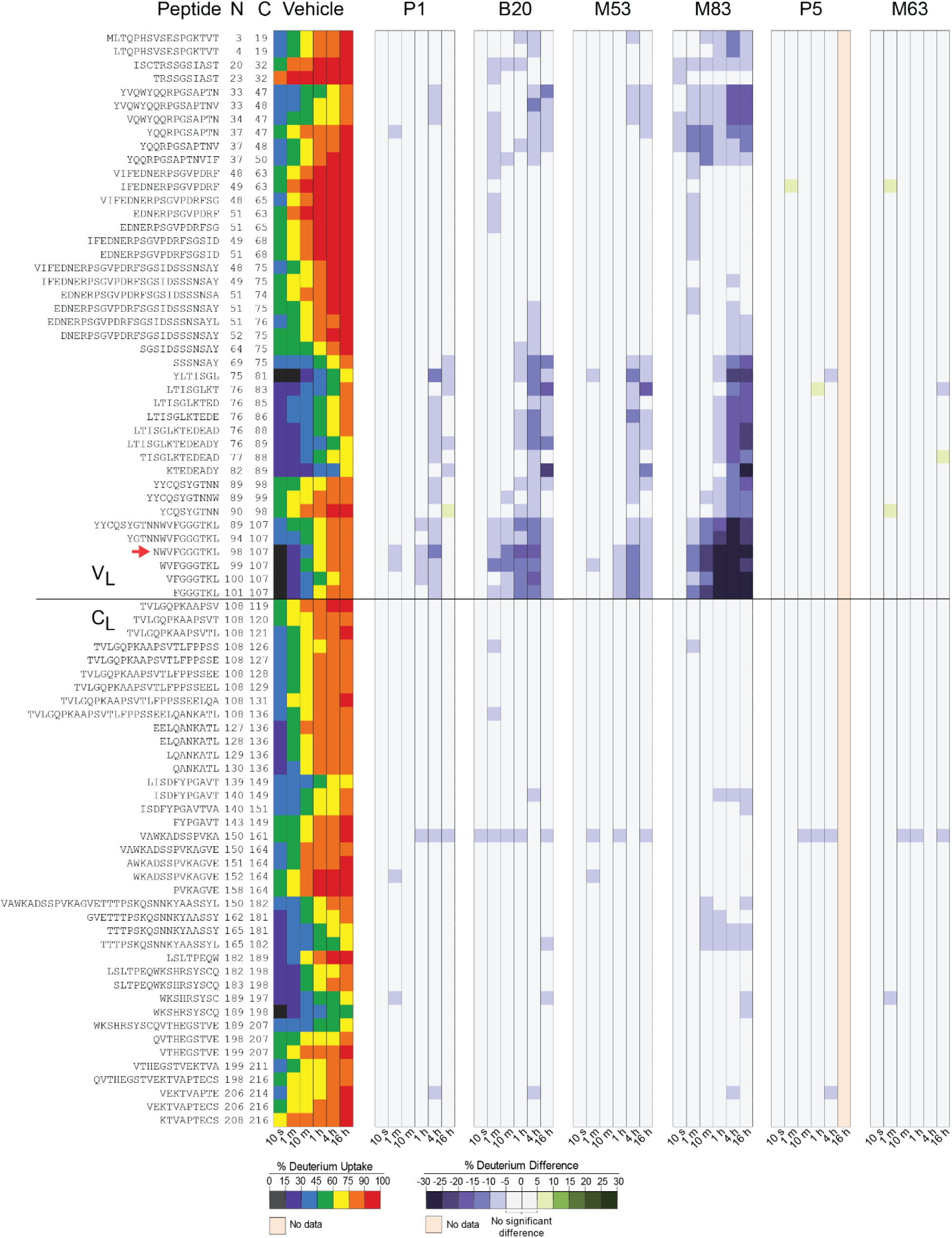
HDX-MS screening of six different kinetic stabilizers. Peptide chiclet plots display differences in the percent deuterium uptake across the sequence of AL LC upon binding of each compound. The compounds screened were P1, B20, M53, M83, P5, and M63. On the left, the absolute uptake of AL LC (no compound) serves as the baseline for comparison. M83 shows the most significant protection, followed by B20, M53, and P1, with decreasing levels of protection. P5 and M63 do not show notable differences in HDX. The horizontal line separates the variable (V_L_) and constant (C_L_) domains. Regions with no data (peach) or no change (gray) are indicated. The AL LC peptide ^101^NWVFGGGTKL^110^ (corresponding to sequential positions 98-107) is highlighted with a red arrow.

**Figure S5.**
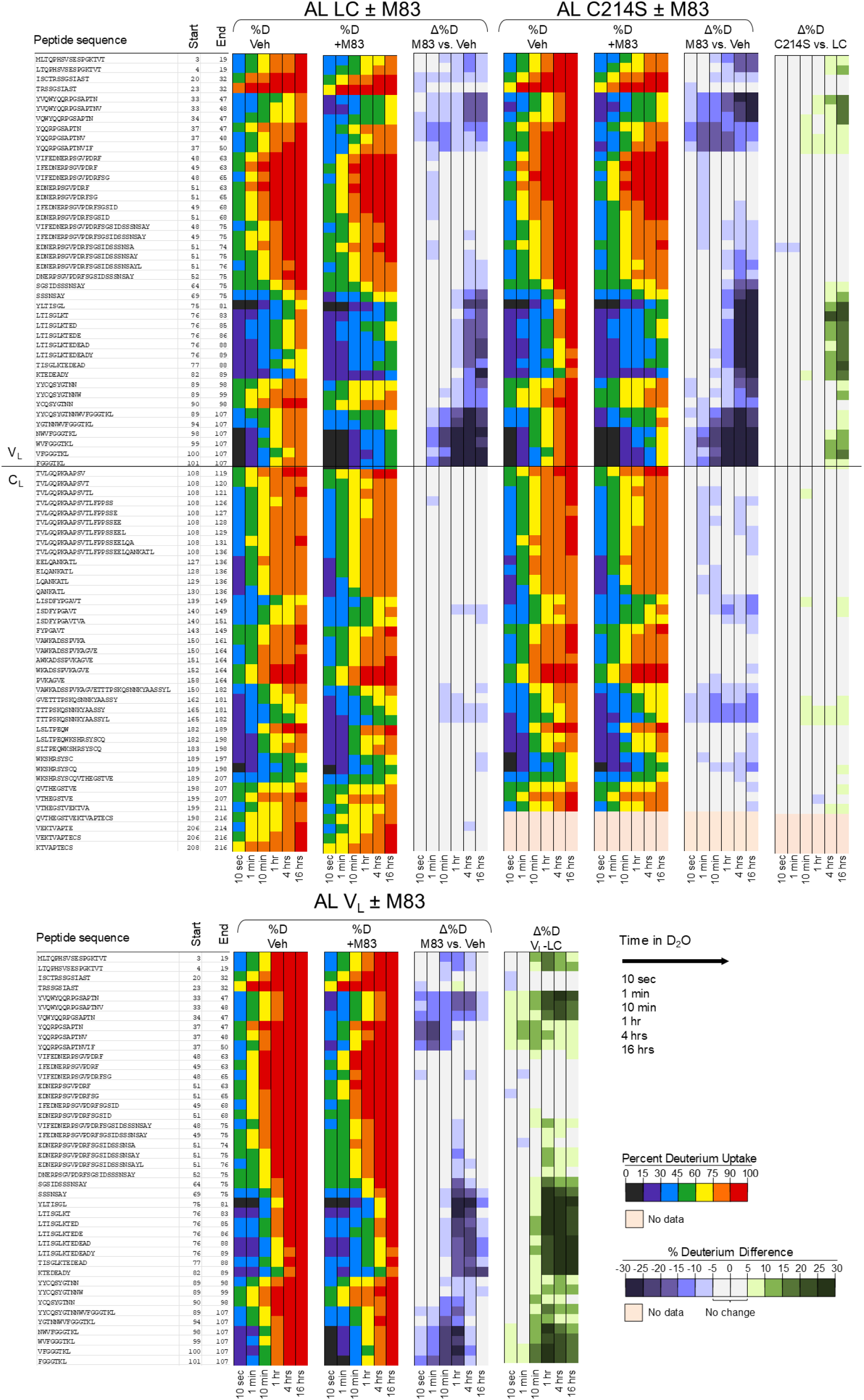
Binding of the kinetic stabilizer M83 to AL LC, C214S, and V_L_ monitored by HDX-MS. This figure presents the same data as Figure 3, displayed in a peptide-resolution chiclet plot format instead of residue-level heatmaps. Percent deuterium incorporation (%D) is shown for the protein in 0.15% DMSO vehicle and in the presence of M83, alongside the differences in deuterium uptake (Δ%D). Comparisons between constructs (C214S-LC and V_L_-LC) are also shown. Horizontal lines separate V_L_ and C_L_ domains. Regions with no data (peach) or no change (gray) are indicated.

**Figure S6.**
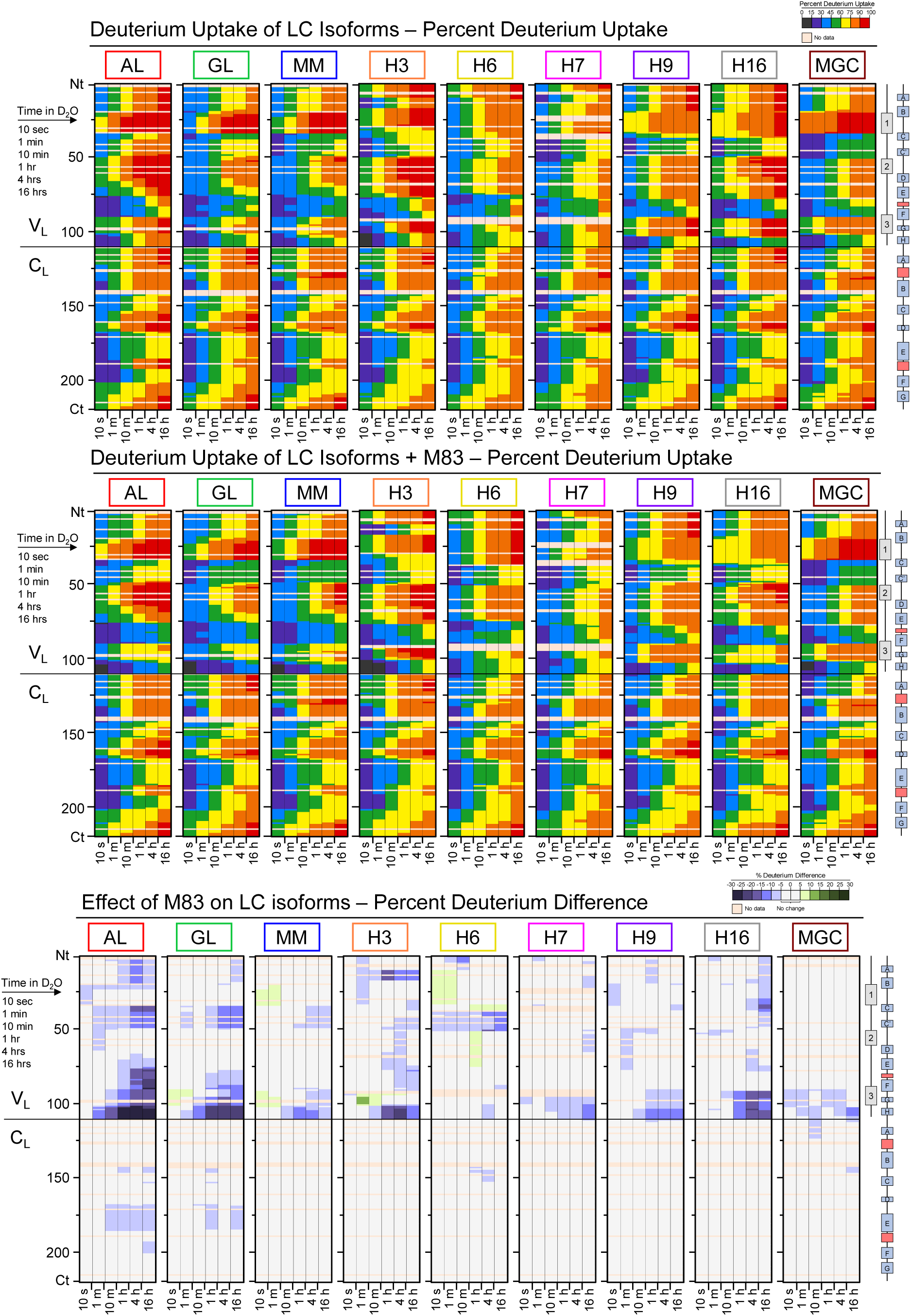
Single residue aligned heatmaps of LC isoforms with and without M83. Top panels: Percent deuterium uptake (%D) for all nine LC isoforms in the absence of M83. Middle panels: Percent deuterium uptake (%D) for LC isoforms in the presence of M83. Levels of deuterium uptake is color-coded as per legend, with red indicating high uptake (low protection) and blue indicating low uptake (high protection). Bottom Panels: Percent deuterium difference (Δ%D) plots showing the effect of M83 on deuterium uptake for each LC isoform. Blue regions indicate areas of increased protection upon M83 binding, while green regions represent areas with reduced protection. The solid horizontal line indicates the boundary between V_L_ and C_L_ domains.

**Figure S7.**
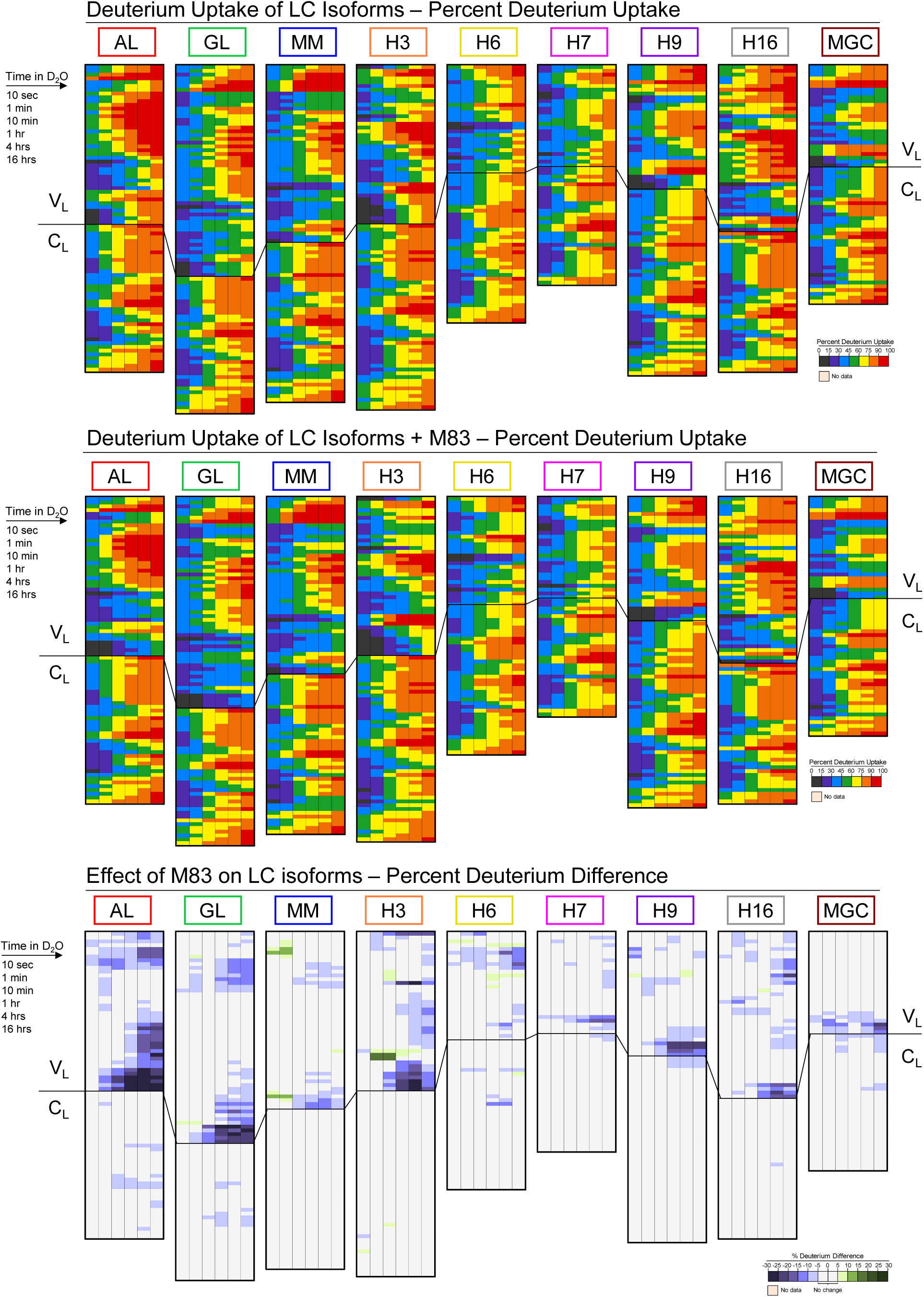
Peptide chiclet plots showing deuterium uptake of various LCs and the effect of M83. Top panel: Percent deuterium uptake (%D) for all nine LC isoforms in the absence of M83. Levels of deuterium uptake are color-coded as per the legend, with red indicating high uptake (low protection) and blue indicating low uptake (high protection). Middle panel: Percent deuterium uptake (%D) for all nine LC isoforms in the presence of M83. Bottom panel: Percent deuterium difference (Δ%D) plots showing the effect of M83 on deuterium uptake for each LC isoform. Blue regions indicate areas of increased protection upon M83 binding, while green regions represent areas with reduced protection. The solid horizontal line indicates the boundary between the variable domain (V_L_) and the constant domain (C_L_).

**Figure S8.**
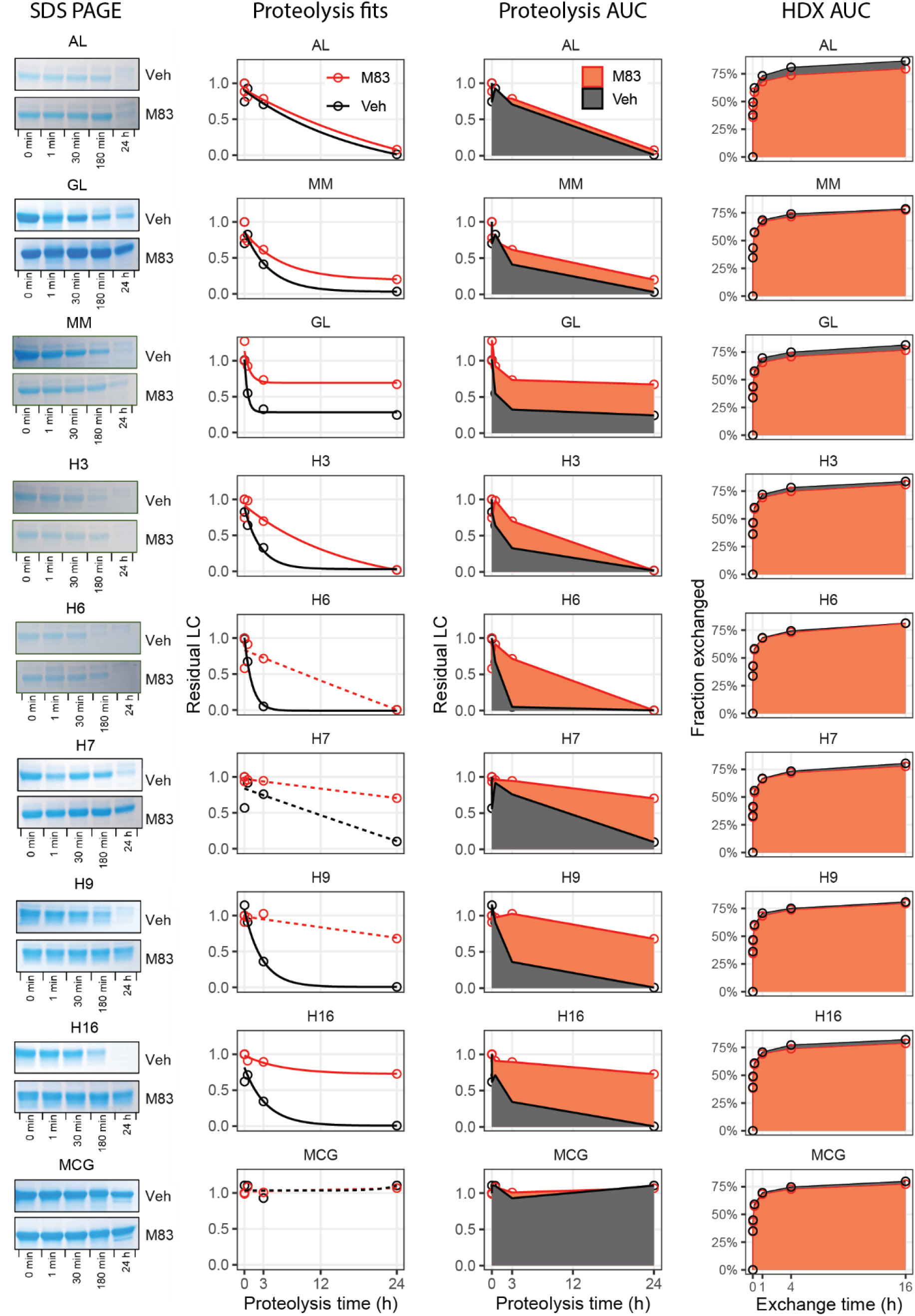
Limited proteolysis assay of FL LCs with and without M83 monitored by SDS-PAGE. FL LCs were tested at a concentration of 8.3 μM, with M83 at 50 μM (1:6 molar ratio) in 10 mM phosphate buffer, pH 7.4, 150 mM NaCl, at 37°C and 450 rpm. Proteolysis reaction was initiated by addition of trypsin at 0.232 μM. Time points (0, 1, 30, 180 minutes, and 24 hours) are shown for digestion experiments. "+" and "-" refer to digestion in the presence and absence of M83, respectively. Gels (10-20%) were run under reducing conditions and stained with Coomassie Blue. Alcohol dehydrogenase was included as a loading control (not shown). Data were fitted to a single exponential decay model and the area under the curve (AUC) was calculated. Dashed lines show data where the fits did not converge. For comparison, the average AUC for all residues was calculated for each FL LC.

